# Cigarette smoke sets up a pro-inflammatory circuit in the lung that induces the hyper-activation of autoreactive T helper cells

**DOI:** 10.1101/2025.08.06.668725

**Authors:** Nuria Alvarez-Sanchez, Annette Haughian, Brendan Cordeiro, Deeva Uthyakumar, Elvira Paneda, Thomas A. Heaney, Molly Pitkethly, Jhenifer Gabriela Vasquez Rojas, Emily Pullen, Elyse Latreille, Chao Wang, Warren L. Lee, Martin Stampfli, Clinton Robbins, Shannon E. Dunn

## Abstract

Cigarette smoke (CS) exposure increases the risk of multiple sclerosis; however, the mechanisms are unclear. To investigate this, we exposed C57BL6/J mice to CS or ambient air (AA) for 8 weeks using a protocol modelling human chronic CS exposure and studied the impact of these exposures on the development of experimental autoimmune encephalomyelitis (EAE). We found that CS increased the incidence of neurological signs in 2D2 myelin oligodendrocyte glycoprotein (MOG) T cell receptor transgenic mice, whereas in active EAE induced by MOG peptide and Complete Freund’s adjuvant, CS delayed the onset of disease. In both cases, the effects of CS tended to be greater in the males who also developed greater leukocyte infiltration and expression of IL-12p40 and other cytokines. To gain insights into the paradoxical effects of CS in these EAE models, we transferred congenically-marked pMOG-reactive T helper cells into AA- or CS-exposed mice and examined the phenotype of these cells in the lungs, spleen, and spinal cord. We found that CS-exposed lungs acted as a sink for the pMOG-reactive T cells, in that more of these T cells homed to the lungs instead of the spinal cord. However, these CS-educated pMOG Th cells exhibited a hyperactivated Th phenotype with greater expression of activation markers, IL-17, and GM-CSF, a cytokine that is essential for EAE development. Intranasal treatment with anti-IL-12p40 negated these effects of CS on the expression of pro-inflammatory cytokines by pMOG Th cells. These studies suggest that CS sets up a pro-inflammatory circuit in the lung that amplifies the encephalitogenic properties of myelin-specific T effector cells.

**Highlights:**

- CS exposure enhances the development of spontaneous neuroimmunity.
- Myelin-reactive T helper 1 responses are enhanced by CS exposure.
- CS-exposed lung attracts myelin-specific T helper cells to this site.
- CS enhances myelin-specific Th1/Th17 cytokine production and GM-CSF through lung IL-12p40

## 1. Introduction

More than a billion people worldwide smoke cigarettes [1]. This habit is a major cause of preventable death [1]. Though tobacco consumption has declined in recent decades [2], consumption rose during the 20th century [3], coinciding with the rising prevalence of autoimmune diseases [4, 5]. In this regard, CS exposure has been linked to an increased incidence of MS, rheumatoid arthritis (RA), systemic lupus erythematosus (SLE), Crohn’s disease, and psoriasis [6]. It has also been estimated that a third of RA cases [7], and 20% of MS cases [8] are preventable if people stopped smoking. For MS, smoking increases disease risk in a dose-dependent manner (odds ratio between 1.4-2.0), with a tendency for a greater effect in males than in females [9]. This effect of CS on the risk of MS is sustained even after 5 years after quitting smoking [9]. In addition, CS hastens disability accrual in people with established MS (pwMS), which is associated with greater disease activity and brain atrophy [10–12]. Each year of smoking post-MS diagnosis is associated with a five percent accelerated time to conversion to secondary progressive MS (SPMS), a stage of disease characterized by a steady accumulation of disability [13]. In contrast, quitting smoking delays onset of SPMS [13, 14]. Despite this strong connection between CS and MS, the biological mechanisms of how CS promotes myelin-specific autoimmunity remain unknown.

Clues to these potential mechanisms have been provided by epidemiology studies that studied the effects of different modes of tobacco consumption on MS risk in humans. While CS exposure promotes MS, consumption of Swedish snuff, a tobacco product that is absorbed through the oral mucosa, protects against this disease [9]. Switching from smoking to snuff also slows the onset of the progressive phase of MS [13]. These data suggest that it isn’t the tobacco components *per se*, but rather the inhalation of combusted smoke particles into the lung that is the risk factor for the disease. Studies have also found an interaction of CS exposure with carriage of the disease-associated MHC-II risk alleles in the development of MS and RA [15–17], suggesting that the smoke particles are somehow promoting the activity of autoreactive T helper cells in these diseases. Consistent with this idea, a study that examined the phenotype of polyclonal T cells in the broncheolar lavage (BAL) of smokers and non-smokers with and without MS noted that T cells showed a hyperactive phenotype in the BAL of smokers [18]. In addition, SLE and RA, smoking promotes autoantibody generation [19–21]. In experimental models of RA, it is also reported that CS promotes protein citrullination in the lung, a post-translational modification where arginine is converted to citrulline [22]. Citrullinated proteins generate cryptic epitopes that serve as new autoantigens that stimulate T cells in people with RA [23]. While protein citrullination is also increased in the brains of people with MS (pwMS), this is not associated with increased T cell reactivity to citrullinated antigens [24], suggesting that CS promotes MS-related autoimmunity by another mechanism. Altogether, mounting evidence suggests that lung pathology caused by the inhalation of tobacco smoke particles promotes T cell autoimmunity; however, these mechanisms have not yet been elucidated.

To gain insights into these mechanisms in MS, we passively exposed male and female mice to CS for 8 weeks; a regimen previously reported to model chronic CS exposure in humans [25]. We then studied the impact of CS on myelin-specific autoimmunity in active (MOG p35-55/CFA-induced) and spontaneous (2D2 MOG T cell receptor transgenic mice) EAE models of MS. We found that CS exposure enhanced leukocyte infiltration and inflammation in the lung and that this was associated with greater IFN-γ production by myelin-specific T helper cells in the peripheral lymphoid organs in the active model of EAE; however, EAE onset was puzzlingly delayed in the male mice and was not affected in the female mice. In contrast, in 2D2 MOG-specific T cell receptor transgenic mice, CS promoted the development spontaneous neuroimmunity characterized by leukocyte infiltration in the dorsal root ganglion and spinal nerve roots. To gain insights into these paradoxical findings, we tracked congenically-marked pMOG-reactive Th17 cells after transfer into CS- or AA-exposed mice. We found in both males and females that CS exposure caused a greater number of the pMOG-reactive Th17 cells to migrate to the lung, which associated with a delayed onset of EAE. In the CS-exposed lungs, the pMOG-reactive Th cells acquired a hyper-activated phenotype and greater expression of IL-17 and GM-CSF. These pMOG-reactive Th cells in CS-exposed retained this greater expression of IL-17 and GM-CSF after reaching the CNS. Lung DCs from CS-exposed mice also had a greater capacity to activate and promote cytokine production by 2D2 cells *in vitro*, an effect that was fully- or partially-prevented by neutralizing IL-12p40. Together, these studies suggest a mechanism whereby CS promotes autoimmunity by recruiting antigen-experienced autoreactive Th cells into the lung, where these cells are activated by a mechanism of bystander activation.

## 2. Results

### 2.1 Chronic CS exposure induces lung inflammation

To investigate the mechanisms linking CS exposure to the promotion of myelin-specific autoimmunity, we employed a whole-body CS exposure system and 8-week CS exposure protocol that has been previously described to model the lung pathology and inflammatory signatures seen in human smokers [25, 26]. Histological studies conducted on lungs from males exposed to CS- confirmed the presence of greater leukocyte infiltration in CS- versus AA-exposed mice (Fig. S1A-C). CD45^+^ leukocytes were detected throughout the parenchyma and within immune microclusters surrounding the bronchioles and blood vessels (Fig. S1A-C). To gain insights into the composition of lung immune cell infiltrates with CS, we also conducted flow cytometry analysis on lung mononuclear cells (Fig. S1D-G). This analysis confirmed greater CD45^+^ leukocyte infiltration in the lungs of males with CS-exposure (Fig. S1D-E). The numbers of eosinophils and conventional DCs were increased in both sexes with CS, whereas T and B cells, alveolar and interstitial macrophages were significantly increased only in the males (Fig. 1G). We also immunolocalized lymphocytes in lung sections in male CS- and AA-exposed mice. We found that B cells were rare, and T cells were sparsely scattered throughout the lungs of AA-exposed mice. In contrast, in CS-exposed mice, T cells were detected in para-bronchiole immune micro-clusters, sometimes in close-proximity to B cells (Fig. S2). Taken together, these data suggested that this CS-protocol enhanced lung inflammation, especially in the male mice.

**Figure 1.**
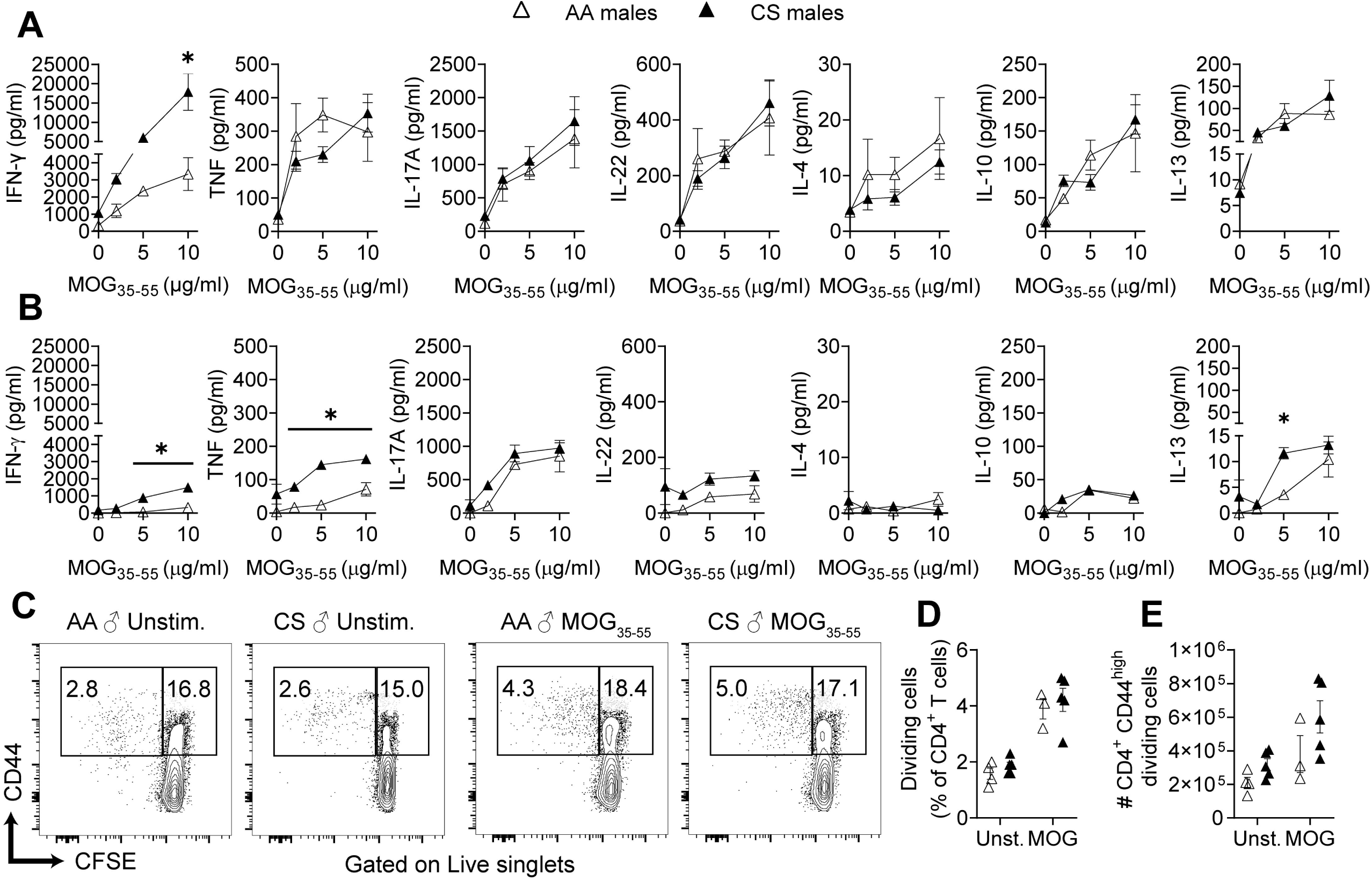
Chronic exposure of mice to cigarette smoke (CS) enhances MOGp35-55-specific Th1 responses. Ambient air (AA)- or CS-exposed male C57BL6/J mice were immunized with MOG p35-55/CFA and the recall cytokine and proliferative responses of cells were assessed 9 days later in spleen and draining lymph node (dLN) mononculear cells. (A-B) Levels of cytokines in supernatants of MOG_35-55_-stimulated cell cultures from spleens (A) and dLNs (B) measured using a multiplex assay. (C-E) Division of splenic CD44^high^CD4^+^ T cells after stimulation in the presence/absence of MOG p35-55 *ex vivo* as assessed by CFSE dilution. Representative plots of dividing CD4^+^ T cells (C), and frequency (D) and total numbers (E) of dividing CD44^high^CD4^+^ T cells. Data in A-B are mean ± SEM of triplicate wells of cultures from pooled mice and representative of three independent experiments in males and two independent experiments in females. Data points in C are individual mice from one experiment representative of 3 proliferation assays performed. *: p≤0.05 between groups as determined by Mann-Whitney (A,B) or 2-way ANOVA and Bonferroni post hoc test (C,D).

### 2.2 Chronic CS exposure enhances myelin-specific Th1 responses and the incidence of spontaneous autoimmunity in 2D2 mice, but delays the onset of active EAE in males

We next examined the effect of this CS exposure regimen on myelin-specific Th cell responses during EAE induced by immunization with MOG_35-55_/Complete Freund’s adjuvant (CFA). Mononuclear cells were prepared from the draining lymph nodes and spleens of mice at 9 days post-immunization, and the MOG_35-55_-elicited cytokine and proliferation response of these cells was examined *ex vivo*. We observed that CS induced a prominent increase in the pMOG-elicited IFN-γ secretion by spleen and lymph node cells of male mice as compared to AA-exposed controls (Fig. 1A-B). pMOG-elicited TNF production was also greater in the spleen with CS, and IL-13 was elevated in the lymph nodes (Fig. 1A-B). In contrast, the levels of IL-17 and other Th2 cytokines were not significantly altered (Fig. 1A-B). The increased pro-inflammatory MOG_35-55_-Th1 response in the lymphoid organs did not relate to there being a greater abundance of MOG_35-55_-responding CD4^+^ T cells (Fig. 1C-E). Greater MOG_35-55_-specific Th1 cytokine production was also seen in the spleen, but not in lymph nodes, when similar studies were done in female CC- and AA-exposed mice (Supplementary Fig. 3A-B). We next repeated these studies but also injected mice with pertussis toxin and followed mice for the development of clinical signs of EAE (Fig. 2A). Since Th1 cells can be pathogenic in EAE [27], we had anticipated that EAE would be more severe with CS exposure. Instead, this treatment delayed and lessened the severity of EAE in the males and had no significant effect on clinical scores in the females (Fig. 2B-E). Histological studies conducted at end-point confirmed lessened CNS inflammation and demyelination in CS-exposed male mice (Fig. 2F-G). This effect of CS in the males was seen regardless of whether CS exposures were continued (Fig. 2A) or stopped prior to EAE induction (Supplementary Fig. 4).

**Figure 2.**
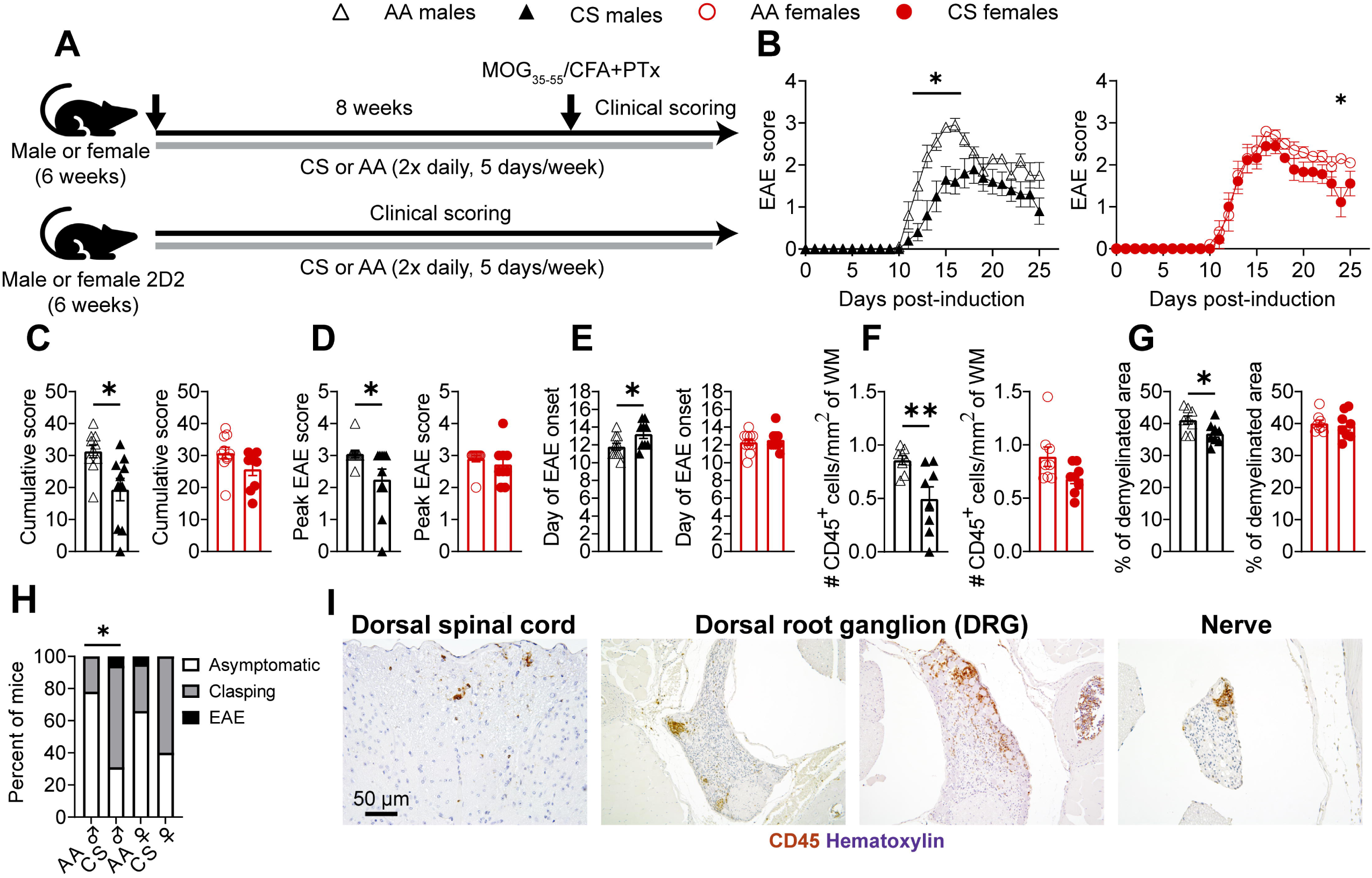
Cigarette smoke (CS) exposure in males decreases the severity of active EAE, yet promotes spontaneous autoimmunity in MOG T cell receptor transgenic mice. (A) Schematic of experiments: C57BL/6J mice were exposed to 8-10 weeks of ambient air (AA) or CS prior to active EAE induction by immunization with pMOG/CFA plus pertussis toxin. Alternatively, 2D2 mice were exposed to AA or CS for up to 8 weeks or until clinical symptoms developed. Black denotes males and red denotes females. (B) EAE clinical scores over time, (C) Cumulative EAE scores, (D) Peak EAE scores, and (E) Day of EAE onset. (F-G) Histological scoring of EAE was done at endpoint by counting the density of CD45^+^ cells/mm^2^ of white matter (WM) (F) or percent demyelination (G) at the level of the thoracic spine in AA- or CS-exposed EAE mice. (H) Frequency of 2D2 mice developing CNS autoimmunity after AA- or CS-exposure. (I) Representative CD45 staining showing sparse immune cell infiltration in the spinal cord, the dorsal root ganglion and spinal nerve roots of CS-exposed 2D2 mice. Clinical data in (H) are from two consecutive cohorts each with (n=9-21/group/sex). *: p≤0.05 between groups as determined by two-way ANOVA and Bonferroni post hoc test (B), by two-tailed Mann-Whitney U test (B-G), or by χ2 test (H).

To investigate whether CS could have an impact on the incidence of EAE, we also tested the effect of CS- and AA-exposures in 2D2 MOG-specific T cell receptor (TCR) transgenic mice (Fig. 2A), which can develop spontaneous EAE when housed under specific-pathogen-free conditions [28, 29]. We found that classic EAE signs rarely developed in either AA- or CS-exposed mice; however, with CS, male mice exhibited an greater incidence of hindlimb clasping upon tail elevation (Fig. 2H). Previously, we reported that this neurological reflex develops in 2D2 mice in response to mild T cell infiltration in the spinal nerve roots and dorsal spinal cord [30]. To confirm this, we harvested the spinal columns and spinal cord from representative mice that developed clinical signs and from mice that remained asymptomatic, and embedded these in paraffin, and stained spinal cord/column sections for CD45. We observed only a small number of CD45^+^ cells in the dorsal white matter and spinal nerve roots of the CS-exposed 2D2 mice (Fig. 2I). The more prominent pathological finding was that 50% of dorsal root ganglions (DRGs) of hindlimb clasping CS-exposed 2D2 mice were infiltrated with CD45^+^ cells (Fig. 2I). In AA-exposed mice, no infiltrating cells were observed in the spinal cord or nerve roots and a significantly lower fraction of DRGs were infiltrated (12%, Chi-square=38.1, p<0.05). Since the DRG does not express MOG and is part of the peripheral nervous system, we suspect that this clinical phenotype is driven by autoimmunity against neurofilament-medium, a protein that is strongly cross-recognized by the 2D2 TCR CD4^+^ T cells [30, 31]: Therefore, the signs we detected likely represents neuroautoimmunity. Altogether, CS enhanced Th1 cytokine production by pMOG-reactive T cells upon pMOG/CFA immunization and increased the incidence of spontaneous neuroimmunity, but paradoxically delayed the onset of active EAE. In both cases, the effects were more prominent in the male mice.

### 2.3. CS attracts myelin-specific Th Cells to the lung where they acquire increased pro-inflammatory characteristics

The results in the active EAE model suggested that the MOG_35-55_-reactive T cells were being primed appropriately, but that CS was somehow preventing their trafficking to the CNS. Past studies demonstrated that myelin-reactive Th cells travel through the bloodstream to the lung and bronchiole-associated lymphoid (BALT), where they reside for several days before continuing to the CNS [32]. To see whether the myelin-reactive Th cells were homing to a greater extent to the lungs with CS-exposure, we employed a passive transfer model of EAE where we transferred CD45.1 congenic female MOG p35-55-reactive cells into male CS- or AA-exposed hosts. The T cells were polarized with IL-23, which is necessary to make these cells encephalitogenic by promoting a pathogenic Th17 phenotype. Similar to findings in active EAE, CS-exposed mice initially showed a delay in disease onset; however, eventually, EAE scores became equivalent between the two groups (Fig. 3A). Flow cytometry on CNS mononuclear cells performed at endpoint revealed a similar accumulation of total leukocytes in CS- and AA-exposed mice (Fig. 3B shows representative gating, results in Fig. 3C); however, less than half the amount of CD45.1^+^CD4^+^ cells made it to the CNS in the CS- vs. AA-exposed mice (Fig. 3D). When we examined cytokine expression within the transferred CD4^+^ T cells in the CNS, we found that a greater fraction of the cells in the CS group expressed IL-17 or co-produced IL-17 and GM-CSF (Fig. 3E-F). This result suggests that the T cells became more highly encephalitogenic in the CS host environment.

**Figure 3.**
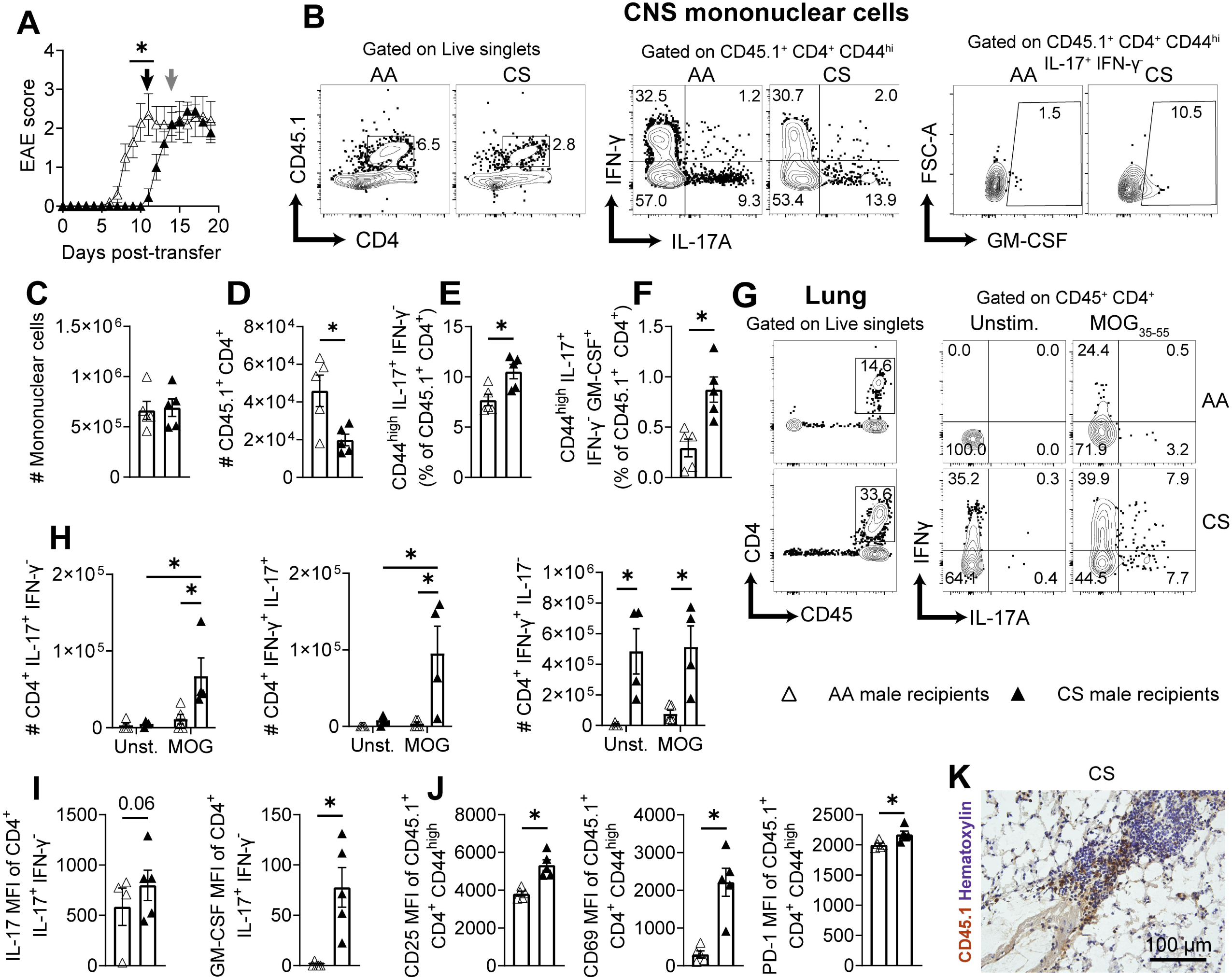
Myelin-reactive T cells are stalled and hyperactivated in cigarette smoke (CS)-exposed lungs. Passive EAE was induced in male C57BL/6J mice that had been exposed to ambient air (AA) or CS, by transferring IL-23-polarized, MOG_35-55_-reactive female T cells from unexposed donor mice. (A) EAE scores in AA- and CS-exposed recipients. (B) Representative flow cytometry plots of CD45.1^+^ T helper cells in the CNS compartment in established EAE (*grey arrows* in A). (C-D) Total numbers of mononuclear cells (C) and of CD45.1^+^CD4^+^ T cells (D); (E-F) frequencies of MOG_35-55_-reactive CD45.1^+^CD4^+^CD44^high^ T cells producing IL-17 (E) or co-producing IL17 and GM-CSF (F). (G) Shows representative FACs plots of CD4^+^ T cells in the lung. (H) Total numbers of IL17^+^, IL17^+^IFN-γ^+^, and IFN-γ^+^ CD4^+^ T cells in lungs. (I) IL-17 and GM-CSF median fluorescence intensities (MFIs) of lung MOG_35-55_–reactive IL17^+^IFN-γ^-^ CD4^+^ T cells. (J) CD25, CD69 and PD-1 expression by donor CD45.1^+^CD4^+^CD44^high^ T cells in the lungs. (K) Immunolabeling of donor CD45.1^+^ cells in the lungs. Data in A are presented as mean ± SEM of individual mice from one representative experiment of at least four performed in males. Data in C-F are mean ± SEM of individual mice from one experiment of two that were performed, one conducted at disease onset and one at peak disease. Data investigating cytokines and activation markers in the lung are mean + SEM of individual mice and are representative of 4 independent experiments. *: p≤0.05 between groups as determined by two-way ANOVA and Bonferroni post hoc test (H) or by two-tailed Mann-Whitney U test (A,C-F, I-J).

To investigate the cause of the delayed passive transfer EAE in the males, we repeated experiments, except harvested the lungs and spleens from mice at a time when clinical signs were delayed in the CS group (e.g. *black arrow*, in Fig. 3A). These experiments revealed a 5-10-fold higher number of MOG_35-55_-specific IL-17^+^ and IL-17^+^IFN-γ^+^ CD4^+^ T cells in lungs of CS relative to the AA-exposed mice (Fig. 3G-H). In contrast, we detected no differences between groups in the spleen (Fig. S5A). In addition, we detected an overall increase in IFN-γ^+^ by polyclonal CD4^+^ T cells in the lung, in the absence of pMOG, consistent with the idea that CS-exposed lung environment has Th1 promoting activities (Fig. 3I-J). In light of past reports that CS exposure can enhance the activation of T cells in the BAL in humans and mice [18, 33], we also measured the T cell expression of activation markers as well as GM-CSF, a cytokine that was upregulated by CS exposure in the CNS-infiltrating Th cells and that is crucial for initiating demyelination in EAE [34]. This analysis revealed greater expression of IL-17 and a massive increase in GM-CSF (Fig. 3I). In addition, we observed the upregulated expression of activation markers CD25, PD-1, CD69 on T cells in the CS-exposed lung (Fig. 3J). In similar experiments performed in females, we also detected a greater recruitment of CD45.1^+^ CD4^+^ T cells to the lung, and these cells also expressed greater CD25 and CD69 expression with CS (Fig. S5B-C); however, the major difference was that the onset of EAE was delayed only by 1-2 d (Fig. S5D). In contrast to findings for the pMOG-reactive Th cells, the expression of activation markers was not altered on the host CD45.2 CD4^+^ T cells in the lung (Figure S5E). Thus, while the effects of CS in increasing T cell cytokine production were seen for both the host and myelin-reactive Th cells, only the myelin-reactive Th cells, which had recently been primed, showed greater expression of activation markers. To investigate whether the myelin-reactive CD45.1 cells homed to the lymphocyte clusters in the lung, we also reserved a few lungs from the experiment in males for CD45.1 immunostaining. We detected CD45.1^+^ cells throughout the lung parenchyma, but also within the para-bronchiole immune clusters that are enriched in the CS-exposed lungs (Fig. 3K). Thus, CS-exposure attracted the MOG p35-55-reactive Th effector cells into the lungs, where these cells became increasingly activated and expressed greater GM-CSF. At the same time, this chemotactic pull of the CS-exposed lung on the T cells seemed to slow their entry into the CNS.

### 2.4. CS exposure amplifies cytokine production and activation marker expression of antigen-activated, but not naive T cells

The increased expression of activation markers on the myelin-reactive T cells in the CS-exposed lung, but not in the spleen, suggested that these T cells were either seeing an antigen that they recognized in the lung or were being activated in an antigen-independent manner via a process of bystander activation. To gain insights into this, we also phenotyped the transgenic CD4^+^ T cells in the 2D2 mice that were exposed to CS or AA for 8 weeks. As was the case in passive EAE, CS increased the accumulation of total leukocytes and tended to increase the numbers of 2D2 CD4^+^ T cells in the lungs (Fig. S6A-B); however, in contrast to the situation in passive EAE, the majority of these 2D2 Th cells had a naive status and the expression of activation markers was not increased on the 2D2 cells in the CS lungs (Fig. 6CD-E). This argues against the smoke particles serving as an antigen for the Th cells. However, we did detect that the expressions of IFN-γ and IL-17A as assessed by MFI tended to be increased in the CD44^hi^ (memory) but not the CD44^lo^ (naive) fraction of lung 2D2 CD4^+^ T cells. These lung-isolated 2D2 cells also produced greater levels of cytokines upon *ex vivo* stimulation with MOG p35-55 (Supplementary Fig. 6F-G). These effects of CS were not observed in the spleen (Fig. S6G-L), confirming that these effects of CS were mediated through the lung. Collectively, these findings suggest that the CS-induced inflammatory environment is attracting myelin-reactive Th cells to the lung where the pro-inflammatory milieu is inducing bystander activation of antigen-experienced Th cells. In the absence of antigen, the CS lung promotes pro-inflammatory cytokine production by the Th cells, but does not increase their activation state.

### 2.5. Alterations in human and murine lung inflammatory signatures with CS exposure

To gain further insights into what inflammatory pathways may be upregulated in the lung with smoking, we conducted an exploratory Gene ontology (GO) enrichment analysis on an existing data set of differentially-expressed genes (DEGs) that were found to be upregulated with CS in the lungs in humans [35]. GO analysis revealed the upregulation of a number of immune pathways with smoking including “regulation of the innate immune response”, “toll-like receptor signaling”, “antigen processing and presentation”, “chemotaxis”, “leukocyte activation in immune response”, and the “initiation of cytokine programs priming the adaptive immune response” (Fig. 4A and Supplementary Table 1). In addition to these top GO biological processes, specific cytokine pathways were enriched with CS, including “TNF”, “IL-6”, “type II IFN”, “IL-12”, and “IL-23” production (Figure 4A, Supplementary Table 1). To validate these findings, we also performed Molecular Signatures Database (MSigDB) Hallmark and Reactome enrichment on the same set of DEGs, with top immune pathways displayed (Figure 4B). The top 8 immune response pathways were similar to those identified in the GO analysis and encompassed innate immune activation, antigen presentation, and molecular signalling programs that are relevant to T cell programming. Leading-edge genes from these enriched pathways, such as *TLR4, TLR8, IRF5, CD86*, and *CXCL1*, overlapped with those in curated cytokine pathways (e.g., IL-12/IL-23) (see Supplementary Table 1), suggesting upregulation of cytokine pathways by TLRs despite weak enrichment of cytokines themselves within the DEGs.

**Figure 4.**
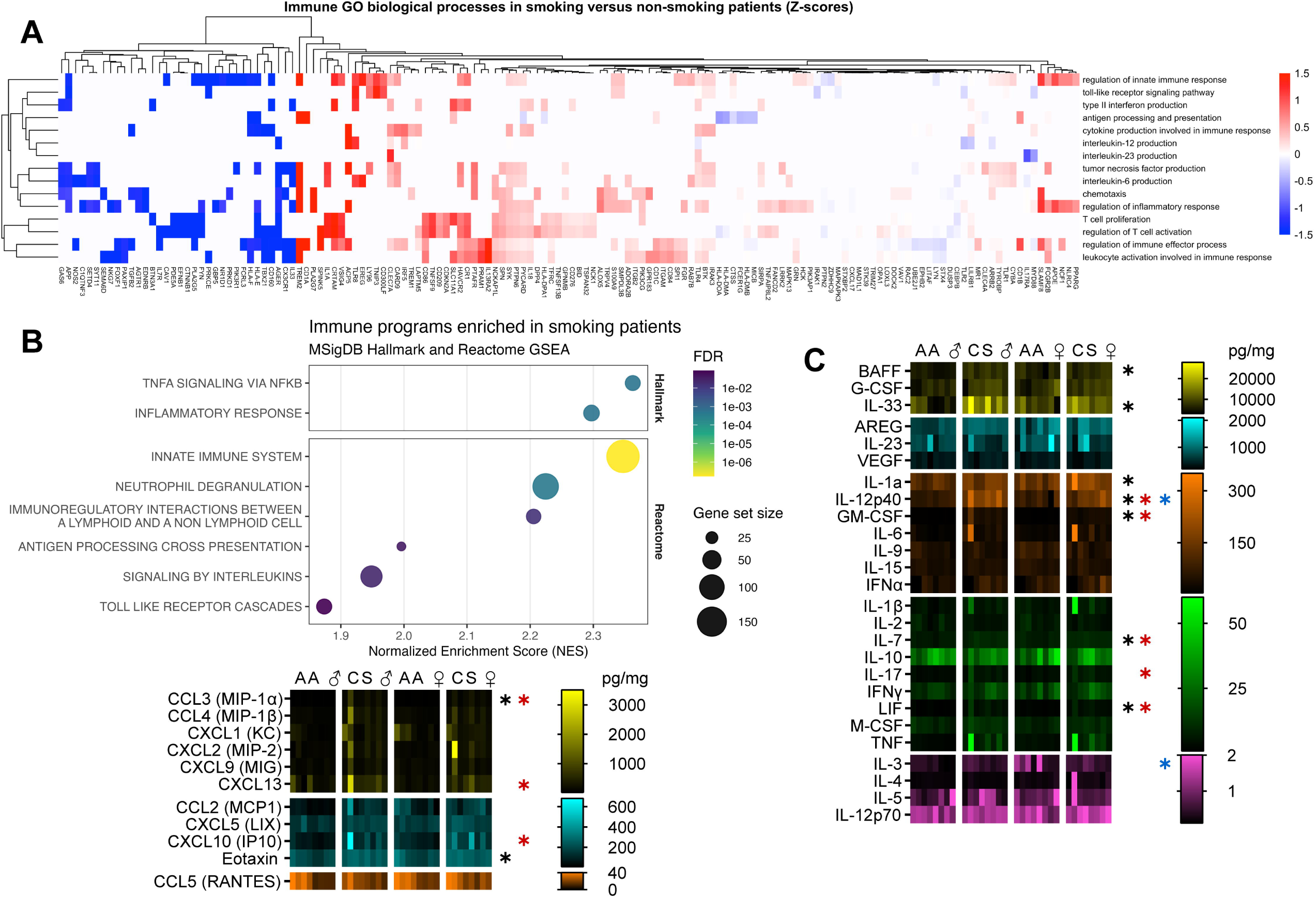
Cigarette smoke (CS) increases pro-inflammatory immune pathways in human lungs and cytokine and chemokine levels in murine lungs. (A) GO term enrichment analysis of microarray-derived differentially expressed genes (DEGs) from human lung tissue of smoking versus non-smoking patients [35]. Significantly enriched immune pathways, and their associated DEGs, were identified using a hypergeometric test with Benjamini–Hochberg correction (FDR > 0.05), with representative immune response and specific cytokine signaling pathways displayed on the heatmap. B) Dotplot of immune response-related MSigDB Hallmark and Reactome gene sets significantly enriched in an exploratory list of microarray-derived DEGs from the human lung tissue of smoking versus non-smoking patients [35]. Significance was determined using the Monte-Carlo method with Benjamini–Hochberg correction (FDR > 0.05). (C) Lysates were prepared from the lungs for AA- or CS-exposed mice that had been immunized with pMOG/CFA. Lung were harvested at d9 post-immunization, were snap frozen in liquid nitrogen, and lungs were homogenized in TPER buffer. Analytes were measured using a multiplex assay and individual ELISA kits. Shown is a heatmap of pg/ml levels of each analyte normalized for total lung protein measured. Black asterisks indicate a difference of CS-group from AA-group in males. Red asterisks indicate a difference of the CS-group from the AA-group in females. Blue asterisks indicate an interaction. Data were analyzed by 2-way ANOVA followed by a Bonferroni post-hoc test. Values represent tissues collected from individual mice from 2 experiments.

To see how inflammatory signatures with our murine CS exposure regimen compared to these human lung gene signatures, we profiled cytokine and chemokine protein expression in lung lysates prepared from AA- and CS-exposed male and female mice (Fig. 4C). Consistent with findings in this human data set and past studies using this CS-exposure regimen in BALB/C mice [33, 36–40], we detected greater levels of several pro-inflammatory cytokines in the lung with CS, including IL-1α, IL-12p40, IL-33, leukemia inhibitory factor (LIF), IL-7, and GM-CSF (Fig. 4C). In contrast, IL-4, IL-5, IL-12p70, IL-23, and IL-10 were not altered (Fig. 4C) . Similar to findings in the human smokers, we detected a trend for increased TNF and IL-6 with CS-exposure (Fig. 4C). Levels of chemoattractants for B cells (CXCL13), eosinophils (eotaxin), monocytes/macrophages (CCL3), and T cells (CXCL10) were also increased with CS (Fig. 4C). When comparing the effects of CS in males and females, we found that certain inflammatory factors were upregulated more so in the males; these factors included IL-12p40, IL-33, eotaxin, and the B cell survival factor BAFF (Fig. 4C). The expressions of IL-3 were also more extensively downregulated in the males with CS (Fig. 4C). The greater levels of eotaxin and BAFF could provide an explanation for the greater eosinophil and B cell with CS in males, whereas the greater expression of IL-12p40 in the CS-exposed mice could explain the greater propensity of the T cells to secrete IFN-γ, IL-17 and GM-CSF [41]. Though IL-33 is known to activate type 2 innate lymphoid cells (ILC2s) in the lung under conditions of allergy, in the presence of IL-12, IL-33 can promote a type 1 inflammatory axis in the lungs [42]. Therefore, we suspect that this could potentially be a further contributor to the greater IFN-γ production by the T cells in CS-exposed lungs.

### 2.6. CS enhances the maturation status of antigen presenting cells in the lung and these are more capable of priming myelin-specific T helper cells

Given the upregulation of the “antigen processing and presentation” pathway, and the prominent increase in the expression of CD86 in the CS-enriched DEGs [35], which also featured in “lymphocyte activation” and “lymphocyte proliferation” pathways, we examined the maturation status of conventional DC (cDCs) and macrophages in the male and female CS- and AA-exposed lungs in the context of passive transfer EAE. We detected that CS increased CD86 expression on lung murine cDCs and macrophages (*black panels*, Fig. 5A). CS also increased MHC Class II expression on lung macrophages and monocytes (Fig. 5A). A similar pattern of greater CD86 expression was seen in the female cDCs and macrophages with CS (Fig. S7). To investigate whether DCs from the CS-exposed had a different functional capacity to prime MOG p35-55-reactive T cells, we magnetically enriched CD11c^+^ cells from lung mononuclear cells from male CS- and AA-exposed mice and co-cultured these cells with male 2D2 CD4^+^ T cells *in vitro* in the presence and absence of MOG p35-55 (Fig. 5B-F). We detected that a greater percentage of 2D2 CD4^+^ T cells were activated, divided, and became effector Th1 and Th17 CD4^+^ T cells when co-cultured with CS lung CD11c^+^ cells as compared to AA-exposed CD11c+ cells (Fig. 5B-F). Notably, MOG p35-55 had to be added in the cultures to see this effect, consistent with the idea that the CS-exposed CD11c^+^ cells were not providing a source of antigen to the 2D2 cells, but rather were enhancing the activation of pre-primed Th effector cells. This suggests that increased co-stimulatory marker expression on lung APCs could be one factor contributing to the greater T cell activation in the CS-lungs.

**Figure 5.**
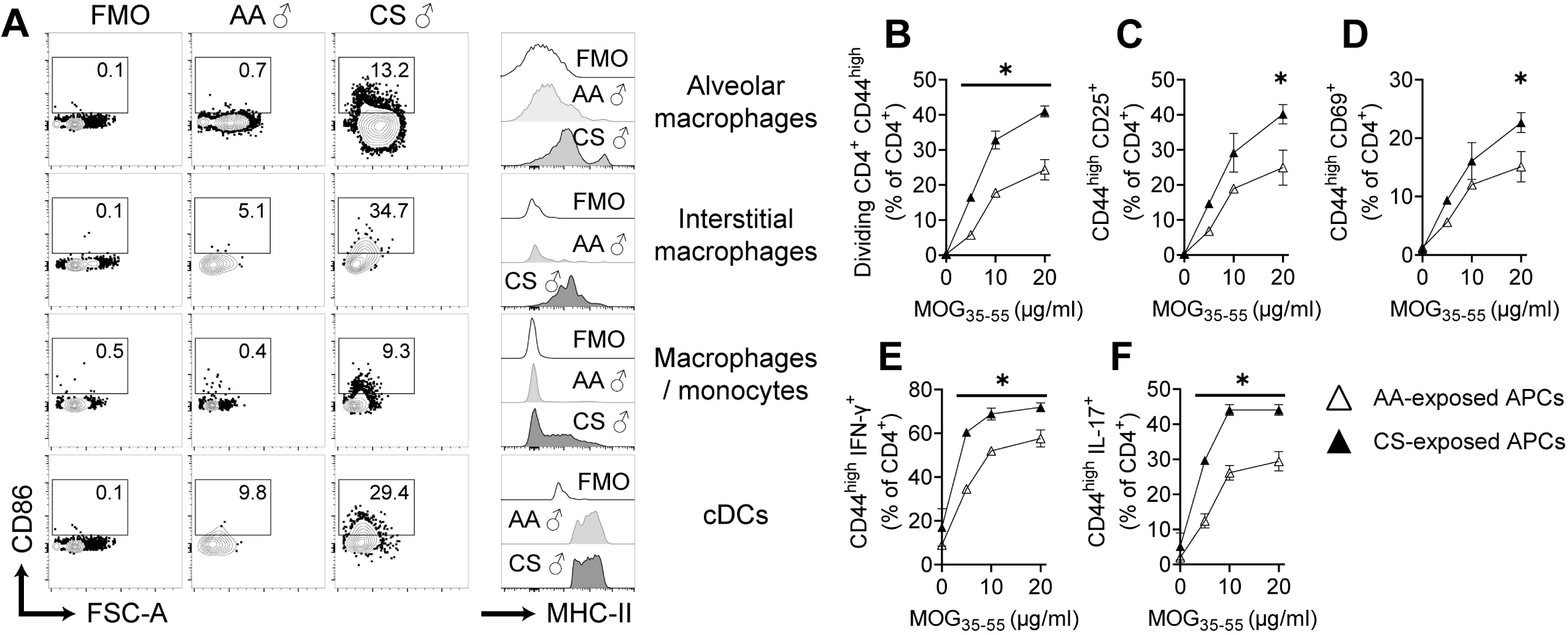
Cigarette smoke (CS) increases the activation of antigen-presenting cells (APCs) in the lung and lung CD11c^+^ cells from CS-exposed mice have a greater capacity to prime myelin-reactive Th cells. Male C57BL/6J mice were exposed to ambient air (AA) or CS for 8-10 weeks and then were transferred with MOG p35-55 reactive T cells. Lungs were harvested at a time when AA-exposed mice had developed EAE and CS-exposed mice were still asymptomatic. (A) Representative FACS plots of the expression of activation markers (CD86, MHC-II) in APCs isolated from male C57BL/6J mice exposed to ambient air (AA) or CS. (B-F) Dendritic cells (DCs) isolated from lungs of AA- and CS-exposed male C57BL/6J mice were cultured in presence of MOG_35-55_ (0-10 µg/mL) with myelin-reactive CD4^+^ T cells isolated from spleens of 2D2 male mice. Proliferation of 2D2 CD4^+^ T cells (B), and expression of CD25 (C), CD69 (D), IFN-γ (E) and IL-17 (F) on memory 2D2 CD4^+^ T cells.

### 2.7. Lung IL-12p40 production mediates the effect of CS exposure in enhancing Th cell cytokine production by the myelin-reactive Th cells

In light of the known effect of IL-12p40 in enhancing Th1/Th17 cytokine production and GM-CSF expression by pMOG Th cells (via IL-12p40 homodimer, IL-23, and IL-12p70) [41, 43], we investigated the role of this cytokine in the effect of CS-exposed DCs on T helper cells. To this end, we repeated the DC: T cell culture studies, except also added anti-IL-12p40 or isotype control antibody to the cultures (Fig. 6A-d). We found that anti-IL-12p40 antibody completely reduced the effect of the CS-DCs in enhancing T cell proliferation (Fig. 6A) and expression of IL-17 (Fig. 6D), and partly inhibited the effect of CS-DC in raising expression of the activation marker CD25 and IFN-γ (Fig. 6B-C), but had little impact in the co-cultures containing AA-exposed DCs (Fig. 6A-D). We also conducted *in vivo* neutralization studies in the context of passive EAE by providing CS- or AA-exposed mice two doses of anti-IL12p40 or isotype control antibody, intranasally on two consecutive days before the transfer of the IL-23-polarized pMOG-reactive Th cells. The phenotype and number of the transferred pMOG-specific Th cells were examined in the lungs at 4 days post-transfer by flow cytometry (Fig. 6E shows representative staining). Pilot studies established that intranasal instillation of IL-12p40 at the 125 µg specifically reduced lung IL-12p40 levels (Supplementary Fig. 8). We found that intranasal administration of this dose of anti-IL-12p40 did not reduce the total leukocyte numbers (Fig. 6F), but did reduce the number of donor pMOG-reactive CD4^+^ T cells (Fig. 6G) and the donor Th17 cells in the CS-exposed lung (Fig. 6G-H). Anti-IL-12p40 treatment also lowered the percentage of CD45.1^+^CD4^+^ T cell cells secreting IL-17 and GM-CSF (Fig. 6I-J), consistent with the idea that IL-12p40 is the major factor accounting for the greater Th1/Th17 effector cytokine production in lung with CS. This treatment, however, did not counter the effect of CS in raising CD25 expression on the T cells (Fig. 6K), suggesting that the T cell activation is mediated by another mechanism, likely increased co-stimulation (Fig. 5). Again, these effects of anti-IL-12p40 were only seen in the CS-exposed lungs (Fig. 6E-K). These findings suggest that IL-12p40 in the lung contributes to both the T cell chemoattractant and the cytokine-promoting properties of CS on the autoreactive Th cells in the lung.

**Figure 6.**
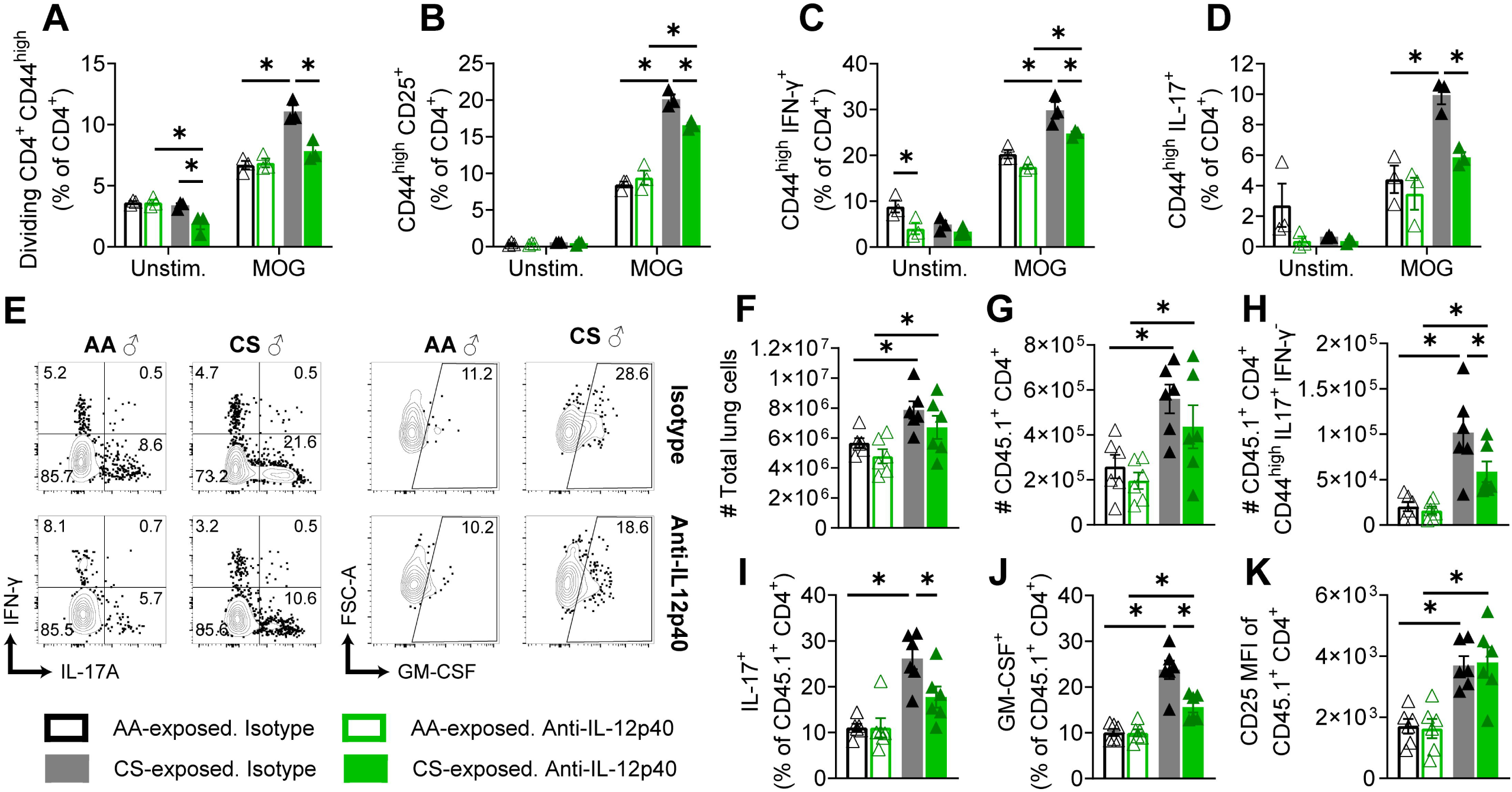
IL-12p40 is a mediator of the effect of CS-exposure in enhancing Th17 and GM-CSF production by myelin-reactive T helper cells. (A-D) AA- and CS-exposed lung DCs were co-cultured with splenic myelin-reactive 2D2 CD4^+^ T cells in presence/absence of MOG_35-55_ (5 µg/mL) with either blocking anti-IL12p40 or isotype control Ab. Proliferation of 2D2 CD4^+^ T cells (A), expression of CD25 (B), IFN-γ (C) and IL-17 (D) in 2D2 CD44^hi^CD4^+^ T cells. (E-K) Passive EAE was induced by transferring IL-23-polarized, MOG_35-55_-reactive T cells from unexposed donor CD45.1^+^ mice into AA-and CS-exposed male C57BL/6J (CD45.2^+^) mice; 2 days prior to, and 1 day after the adoptive transfer. Recipient mice received blocking anti-IL12p40 or isotype control Ab intranasally (125 µg/animal/dose). (E) Representative cytokine staining in the donor CD45.1^+^ CD4^+^ T cells. (F-H) Numbers of total CD45 cells (F), donor CD4^+^ T cells (G) and MOG_35-55_-reactive donor CD44^high^CD4^+^ T cells producing IL-17 (H). (I-J) Frequencies of donor CD4^+^ T cells expressing and IL-17 (I) with GM-CSF (J). (K) Expression of CD25 on the donor CD4^+^ T cells. Data presented are mean ± SEM of triplicates of one experiment (A-D) or are individual mice (F-K). *: p≤0.05 between groups as determined by two-way ANOVA and Bonferroni post hoc test.

## 3. Discussion

The mechanisms that drive MS initiation are unknown. Learning the biology of how MS risk factors enhance autoimmunity may shed light on the pathways involved in the initiation of this disease. Cigarette smoking is known to promote the development of MS and other autoimmune conditions, but no unifying mechanism has been proposed. Here, we demonstrate that CS exposure promotes leukocyte infiltration and the expression of cytokines and chemotactic factors in the lung and induces the maturation of local lung cDCs and macrophages. This has the effect of drawing myelin-specific Th memory cells into the lungs, where these cells express higher levels of cytokines, including GM-CSF, and become hyperactivated. pMOG Th cells that were educated in the CS mouse also showed greater expression of pro-inflammatory cytokines when they reached the CNS compartment. Our studies also revealed that lung IL-12p40 is one mediator of these effects of CS in the lung. Our findings thereby suggest that CS is not the trigger for autoimmunity, but rather amplifies the encephalitogenic potential of myelin-specific T memory cells via a mechanism of bystander activation. In the 2D2 model, we showed that these effects of CS tipped the balance towards autoimmunity.

We explored the effect of CS on EAE development in three EAE models: active EAE, IL-23-induced passive T cell transfer EAE, and in 2D2 mice that can develop spontaneous EAE. In the case of active EAE, where the presence of heat-killed tuberculosis in CFA promotes a strong Th1 response, CS enhanced IFN-γ and TNF production by MOG p35-55 Th cells in the peripheral lymphoid organs. In light of past studies that myelin-reactive Th cells also traffic through the lung during active EAE [32], we speculate that MOG p35-55-reactive Th cells acquired the greater capacity to produce IFN-γ because they had already travelled through the pulmonary circuit at the time of sampling. Greater IFN-γ expression was also observed with CS exposure in the host CD44^hi^CD4^+^ cells that had been recruited to the lung during passive transfer EAE and in the CD44^hi^, but not naive 2D2 TCR transgenic cells in the CS-exposed mice. In the Th17-passive EAE model, CS instead promoted greater T cell expression of IL-17 and GM-CSF by the Th17 effectors, essentially enhancing the expression of cytokines that these cells were programmed to make. Altogether, our findings align with past reports that CS enhances polyclonal T cell activation in the lungs of mice [40, 44] and promotes type I inflammation in the BAL, and IFN-γ^+^IL-17^+^ and CXCR3^+^ (Th1-like) memory T cells in the blood of human smokers [45, 46]. Our finding that CS increased the maturation of murine lung DCs also coincides with past reports that lung DCs play a role in enhancing Th1 and Th17 responses in human smokers with emphysema [47] and in in experimental emphysema induced by long-term CS exposure [48]. In a longer-term, 6 month CS-exposure model, lung DCs also promote the formation of tertiary lymphoid structures in the lung, a pathological feature of the human chronic obstructive pulmonary disease (COPD) [49]. Altogether, these findings suggest that cDCs in the CS-exposed lung amplify cytokine production and promote the bystander activation of the myelin-specific T cells that are attracted to this site.

We also found that neutralizing IL-12p40 in the airways inhibited the recruitment of myelin-specific T cells and reduced the potential of these cells to make Th1 and Th17 cytokines *in vivo*. In the more simplified CD11c^+^: 2D2 T cell co-culture system, anti-IL-12p40 also inhibited the effect of DCs on the MOG-elicited proliferation and activation of 2D2 CD4^+^ T cells. Our proteomics study detected greater IL-12p40, but not IL-12p70 and IL-23 with CS in the murine lung, suggesting that the activities may be mediated through the activity of the IL-12p40 homodimer. Although T cell chemoattractant activities of IL-12p40 homodimer have not yet been reported, this cytokine is chemotactic for both dendritic cells and macrophages (reviewed in [50]) and promotes lung inflammation during viral infection through immune cell recruitment [51]. Previously, we also showed in the CD11c^+^: 2D2 T cell co-culture system that IL-12p40 levels correlated with the expression of both IFN-γ and IL-17 production by 2D2 CD4^+^ T cells and that neutralization of the activity of the IL-12p40 homodimer reduced T cell production of IFN-γ, whereas the IL-17-promoting activities were mediated through IL-23 [43]. Of these IL-12 family cytokines, only IL-12p70 and IL-23 have been reported to promote GM-CSF expression by T cells [52, 53]. Thus, it is likely that a combination of activities of the IL-12p40 homodimer, IL-23 (p19/p40), and possibly IL-12p70 are mediating these effects of CS on cytokine production by T cells. Consistent with this idea, past studies have described roles for both IL-12p70 and IL-23 in the development of other CS-associated lung pathologies, including fibrosis [54] and emphysema [55]. Resolving the individual contributions of IL-12p70, IL-23, and IL-12p40_2_ will require employing mice that are singly or doubly deficient in the protein subunits as has been done in the context of experimental lung infection (e.g, [56]).

Finding that anti-IL-12p40 can dampen myelin-specific Th cell cytokine production aligns with past evidence in EAE that this cytokine plays a key role in the initiation of this disease [57, 58]. Due to promising effects in EAE models, anti-IL12/23 p40 antagonistic antibodies were tested in two phase 2 clinical trials in MS. One trial testing subcutaneous ustekinumab failed to show efficacy of the antibody in reducing the appearance of new brain gadolinium-enhancing lesions in relapsing-remitting MS patients [59], while another trial that tested briakinumab noted an effect in reducing gadolinium-enhancing lesions and disease relapses[60]. This antibody was not developed further since the efficacy was modest compared with other MS treatments and since trials in psoriasis revealed a signal for increased risk of serious infections, nonmelanoma skin cancers, and cardiovascular events [61]. Notably, these trials did not segregate data by sex or smoking status, making it unclear whether anti-IL-12p40 preferentially benefited smokers or males with MS. No one has yet tested the effect of intranasal delivery of anti-IL-12p40 in MS, although other biologics including anti-CD3 are now being tested through this route with promising results (e.g., [62]).

The protein signatures in murine lungs revealed a prominent increase in the alarmins IL-1α and IL-33, IL-12p40, GM-CSF with CS exposure. This cytokine signature is similar to that previously described using the same CS-exposure protocol in mice [36, 37]. These past studies showed that the alarmin IL-1α acts as an upstream effector in the pro-inflammatory cascade in the lung and that neutralizing this cytokine inhibits GM-CSF induction [36], placing IL-1α upstream of GM-CSF in the inflammatory pathway. Additionally, a study focusing on GM-CSF demonstrated that neutralizing this cytokine prevented the accumulation of dendritic cells, the activation of CD8^+^ T cells, and the induction of IL-12p40 in the lungs with CS exposure [37], thus positioning GM-CSF as an intermediary in the pathway leading to IL-12p40 in CS-induced lung inflammation. Our data further revealed that IL-33 was elevated more significantly in male lungs with CS. Although IL-33 is primarily known for it role in promoting expansion of type 2 innate lymphoid cells (ILC2s), especially in female lungs, during states of health and allergy [63], in the context of cigarette smoke and influenza, increased IL-33 coupled with decreased IL-33 receptor ST2 on ILC2s instead promotes macrophage and dendritic cell activation and type 1 immunity [42]. In the context of CS, IL-33-deficient mice are protected from type I inflammation in the lung [42]. Similarly, in humans, the combination of IL-12 and IL-1β is reported to induce trans-differentiation of human ILC2s to ILC1s [64]. Consequently, it is likely that IL-33 worked together with IL-12p40, CCL3, and eotaxin to enhance lung inflammation with cigarette smoke in males.

We noted a trend for greater leukocyte infiltration in the lungs of males with CS. This sex difference did not impact the activation status of either the DCs or of the infiltrating pMOG-specific Th cells, since these measures increased to a similar extent in males and females with CS. However, the greater lung inflammation correlated with comparatively greater delay in EAE in the passive EAE model, which is likely due to a delayed exit pMOG-reactive Th cells from the lung and delayed migration to the CNS. By contrast, this same tendency for greater lung inflammation in males favored the development of neuroimmunity in males in the 2D2 model. The disparate effects of CS in these models can be reconciled when considering how pMOG Th cells are primed in each disease model. In the case of active EAE and passive EAE, pMOG Th cells were primed in the draining lymph nodes by their antigen coupled with a strong adjuvant, and these Th cells travel in a large wave to the CNS to initiate disease. The CS lung, though enhancing the encephalitogenic properties of these cells, also slowed their trafficking to the CNS. However, in the case of the 2D2 model, no external source of antigen was provided, and instead, these cells are thought to gain antigen experience upon exposure to gut commensals and microbes at mucosal surfaces [29, 65]. In this case, the CS lung had the effect of drawing these otherwise dormant 2D2 CD44^hi^ CD4^+^ T cells into a pro-inflammatory environment that had the effect of tipping the balance towards autoimmunity. We expect that this model may be closer to the scenario in human MS where autoreactive T cells are also activated by the microbiota and EBV infection at mucosal sites.

The reasons for these sex differences in lung inflammation with CS are unclear. We did not pursue the mechanisms for the greater expression of IL-12p40 or IL-33 in the males with CS. This is because it has already been established that these cytokines are produced more prominently in females due to the action of sex hormones. IL-12p40 is produced at greater levels by murine female DCs and macrophages in response to T cell-derived signals [66–68]. This sex difference has been demonstrated to be due to enhancing effects of estradiol and repressive effects of androgens on IL-12p40 in macrophages and DCs [66, 67]. Similarly, females have been reported to exhibit greater activity of the IL-33/ILC2 axis in the lung at steady state and during allergic inflammation; this sex difference is due to the effects of androgens in suppressing the lung accumulation of ILC2 cells [63, 69, 70]. We therefore speculate that the sex difference in lung production of certain factors is regulates by effects of sex on signaling pathways upstream. In this regard, analysis of the human CS-induced DEGs identified genes involved in TLR signaling (TLR2, TLR4, MYD88) signaling, activation of the inflammasome (PYCARD/IL18) and activation of Dectin-1-fungal sensing pathways (CLEC7, SYK, CARD9) in the lungs with CS (Supplementary Table 1). Past studies in mice established that these same pathways are upregulated in the murine lungs with CS exposure; the activation of TLR4 is mediated by HSP60 [71], while activation of the fungal sensing pathways mediated by TLR2 and Dectin-1 are in response to greater levels of Hmgb1 [72]. Future studies will explore which of these innate pathways are more operative in the male versus female lung with CS exposure.

We observed that chronic CS in mice also promoted the formation of small T cell and T cell/B cell microclusters in the lungs. These are likely a prelude to the more organized tertiary lymphoid structures reported to develop with more prolonged CS exposures leading to COPD-like features in mice [26] and humans [65]. We showed that the CD45.1^+^ MOG p35-55-reactive Th cells that we transferred homed to these clusters, suggesting that these could be sites where the MOG p35-55 T cells becoming activated. Past studies showed that treatment with a BAFF-Fc reagent could prevent the development of tertiary lymphoid structures in the lungs of mice exposed to CS for 6 months [26]. Interestingly, interstitial lung disease with prominent lymphocytic infiltration has been described as a disease feature of RA [73]. Whether this also occurs in people with MS who smoke, to our knowledge, has not yet been examined; however, the risk of MS is reported to be 6-fold higher in those who subsequently develop COPD before age 60 years [74], hinting at a potential link between lung disease and MS.

Our study also brings further attention to the lung-brain axis in the regulation of MS-related autoimmunity. A connection between the lung and CNS was first suspected when epidemiology studies detected a temporal association between respiratory tract infections and MS onset [75] and disease relapses [76–81]. However, a concrete link was not envisaged until it was demonstrated in studies in rats that myelin-reactive T cells traffic through the BALT and lung draining lymph nodes on their way to the spinal cord during passive and active EAE [32]. The transit through these organs was shown to be important for the acquisition of expression of chemokine receptors and adhesion molecules on the T cells that are necessary for transmigration into the CNS [32]. Other studies have demonstrated that lung commensals and microbes also modulate EAE severity. For instance, intranasal infection of mice with PR8 influenza exacerbates active EAE by promoting a Th1 cytokine response by myelin-reactive Th cells and induces immune infiltration into the spinal roots in 2D2 mice, similar to the pathology that we observed with CS [65, 82]. In addition, it has been shown that depleting the lung flora with neomycin in mice ameliorates EAE by lowering the activation state of microglia in the CNS [83]. Finally, a murine study identified the lung as a promising site for the delivery of antigen-coated microparticles to induce tolerance. Interestingly, part of the benefit of antigen nanoparticle therapy in attenuating EAE related to its ability to promote mild inflammation in the lung, which caused the T cells to be retained in the lung instead of going to the brain [84].

### Study Limitations

Despite testing CS in three different EAE models and demonstrating an effect of CS in promoting neuroimmunity, we were unable to model an effect of CS in worsening *myelin*-specific autoimmunity. Having such a model would be important for pre-clinical testing of therapies that aim to treat lung inflammation in pwMS who smoke. Similar to our findings, others similarly showed that CS-exposure delayed EAE [85], and the onset of autoimmunity in the MRL-*lpr/lp*r SLE-prone mouse strain [86]. Interestingly, in the latter model, the CS had the paradoxical effect of enhancing autoantibody generation. Future studies will modulate the dose of CS and timing of CS exposures to minimize this chemotactic activity while still promoting the pro-encephalitogenic activities of the T myelin-reactive cells.

In conclusion, our findings demonstrate that myelin-reactive Th cells traffic to the CS-exposed lung where signals, including IL-12p40 and increase DC co-stimulation, and amplify T cell activation and cytokine production, thereby providing a general mechanism for how CS may promote autoimmunity.

## 4. Methods

### 4.1. Sex as a biological variable

Our study examined both male and female animals, and the phenotypes were compared between both males and females.

### 4.2. Mice

Male and female C57BL6/J mice (Stock#664), female CD45.1 congenic mice on the C57BL6/J background (PepBoy; B6.SJL-Ptprca Pepcb/BoyJ; Stock# 002014), male and female 2D2 TCR transgenic mice (2D2 mice; C57BL/6-Tg(Tcra2D2,Tcrb2D2)1Kuch/J; Stock#6912) were from the Jackson Laboratory. 2D2 mice were bred in pairings of heterozygote males with wild-type C57BL/6J females. 2D2 heterozygote offspring were genotyped by flow cytometry of PBMCs after staining for CD4 and TCR Vβ11 chain antibodies as described previously [43]. All mice were housed within a specific pathogen-free facility at the University Health Network (UHN), and work was conducted under an animal use protocol (AUP 5937) that was approved by the UHN animal care committee, following guidelines of the Canadian Council on Animal Care.

### 4.3. Exposures to CS and AA, and Cotinine Measurements

Adolescent mice were exposed to CS from research-grade cigarettes, with filters removed (Tobacco and Health Research Institute, University of Kentucky, Lexington, KY) using a whole-body exposure system (SIU48; Promech Lab Inc., Vintrie, Sweden) that was housed in a fume hood [33]. CS was puffed into a central chamber where freely moving mice were housed in compartments with cage mates, with food and water removed. Exposures were initiated at 7 weeks of age and consisted of two daily 50 min sessions of exposure to CS (separated by 3 h) generated from 12 serially burned cigarettes, 5 days/week for 8-10 weeks, and was preceded by an initial 3-day acclimatization period where mice were placed in the exposure box for increasing lengths of time. This CS exposure regimen was well tolerated. AA-exposed controls were brought into the same room, and, at the time of CS exposures, placed into an empty cage having similar dimensions as the caging system used for the CS-exposures, with water and food removed. To estimate the dose of acute CS exposures, blood was drawn within 1 h of exposure and was centrifuged (5000 rpm x 5 min) to collect serum, which was cryopreserved at -80°C. Cotinine was measured using an ELISA kit (Origene) and was found to be 379.1 ± 32.2 ng/ml in CS-exposed mice and 0.4 ± 0.1 ng/ml in the ambient mice, which is similar to levels reported in past studies using this CS-exposure protocol [33] and for chronic human smokers [87].

### 4.4. Lung Pathology

Mice were anesthetized with isoflurane and perfused with 1x PBS containing 5 U/mL of heparin. The lung and heart were dissected, and a 0.5 cm length of tubing (VWR 427411), was inserted into the trachea and was tied in place using surgical twine. To prepare formalin-fixed paraffin sections, lungs were first inflated with formalin by filling a 50 mL syringe with neutral buffered formalin, which was gravity-fed through surgical tubing to a 23 G needle placed in the canula. After the lungs were fully inflated, the trachea was tied off below the level of the cannula, and the heart and lung were stored in 10% formalin. After 3-7 days, the lungs were grossed, processed and embedded in paraffin with the right lung embedded longitudinally, and the left lung upper, middle, and lower lobes embedded in cross-section. H&E staining and CD45 IHC (rat anti-CD45 Ab, clone 30F-11, BD) were conducted as previously described using our established protocols [88]. Dual CD3 and B220 IHC was conducted via a similar method with the following modifications. After de-waxing, antigen retrieval, and blocking [88], sections were stained overnight at 4°C using a rabbit polyclonal anti-human CD3 Ab (1:1000, Dako). The following day, sections were washed 3x in 0.1% PBS-Tween-20 (PBST), probed with secondary Ab (HRP-conjugated, goat anti-rabbit Ab, clone BA-1000, Vector), washed 3 times, and then incubated with HRP-streptavidin complexes (universal ABC-HRP kit, Vector labs) [88]. After three final washes, immune complexes were developed using DAB (Vector SK4100) [88]. Sections were then re-blocked in blocking buffer (2% BSA in 2% rabbit serum) that contained avidin (Avidin Biotin kit, Vector labs) and incubated overnight at 4°C with anti-B220 Ab (1:500, clone RA3-6B2, BD Biosciences) in blocking buffer that contained biotin. Slides were washed with PBST and incubated with biotinylated rabbit anti-rat secondary Ab (1:200, Vector labs) for 30 min at RT. After an additional wash, sections were incubated for 30 min at RT with Avidin-Alkaline Phosphatase (1:1000; BioShop) in 1x PBS. Slides were washed again with PBST, and antibody-AP complexes were developed using Vector Blue as a chromogen (Vector Labs). Slides were dehydrated and then cleared with Citrosolve and cover-slipped using Permount (both from Fisher).

For CD45.1 immunohistochemistry, the lungs and heart with attached trachea were harvested and canulated as described above, except that the lungs were inflated with 500 µL of a 2:3 solution of 1x PBS: OCT compound using a 23 G needle and syringe. The inflated lungs were tied off below the cannula and submerged in the PBS:OCT solution until all specimens were collected. After grossing the lungs, they were embedded in OCT in melting 2-methyl butane cooled by liquid nitrogen. OCT-embedded lungs were cut at 4 µm and stained using a CD45.1 Ab (1:1000, clone A20, Biolegend) and the mouse on mouse (M.O.M) kit (Vector Laboratories) according to kit directions. In brief, sections were air-dried for 10 min and then fixed in ice-cold methanol for 15 min. Slides were washed in 0.5% PBST (3 x 5 min) and peroxidase quenched with 3% hydrogen peroxide in methanol. Sections were blocked using the blocking solution provided in the M.O.M. kit for 30 min at RT, then washed with PBST. After secondary Ab detection, sections were incubated with streptavidin-HRP (universal ABC HRP kit, Vector labs) and then developed with DAB, counterstained with hematoxylin, dehydrated, cleared with xylene, and mounted with Permount™ according to our published protocols [88].

### 4.5. Isolation of Lung Mononuclear Cells

The lungs were dissected as above and placed in a conical tube containing ice-cold RPMI 1640 (Life Technologies) and 2% FBS (Wisent). Lung lobes were transferred to a petri dish containing 5 mL of ice-cold RPMI, scissor-minced and transferred to a 50 mL conical tube. Five mL of 2x digestion buffer containing 300 U/mL type 1 collagenase (Worthington) and 10 U/mL DNAse I (Sigma) in RPMI were added and the tissue was agitated for 1 h on a 37°C shaker. The cells were then dissociated through a 70 micron sieve and centrifuged at 300 x g for 10 min at 4°C. The supernatant was discarded and red blood cells were lysed with ACK buffer (0.15 M NH_4_Cl, 10 mM KHCO_3_, 0.1 mM Na_2_EDTA) for 1 min and 15 s [43], and the lysis stopped with RPMI. Cells were centrifuged again, counted and washed with FACS buffer.

### 4.6. Flow Cytometry

Cells were incubated with Fc Block (5 µg/mL; BioLegend) for 15 min at 4°C and washed in FACS buffer. Cells were incubated with the Ab cocktails detailed in Supplementary. Table 2 and either Fixable Viability Dye eFluor 506 or LIVE/DEAD™ Fixable Aqua Dead Cell Stain (both 1:1000; Invitrogen) in the dark for 30 min at 4°C, and then washed again. If intracellular staining was performed, cells were fixed with 4% paraformaldehyde in 1x PBS for 10 min at 4°C, washed, and then incubated with 1x Perm Wash buffer (BD Biosciences) in the dark for 30 min at 4°C. After washing cells three times with Perm Wash buffer, cells were incubated with the intracellular antibodies detailed in Suppl. Table 2 diluted in Perm Wash buffer for 30 min at 4°C in the dark, and washed with 1x Perm Wash buffer. Cells were either analyzed and unmixed using a Sony SP6800 Spectral cell cytometer and software, or analyzed using a Fortessa X-20 or a Cytoflex-LX cytometer, and data analyzed in FlowJo (version 10). Cell populations were gated using fluorescent minus one (FMO) controls.

### 4.7. MOG p35-55/CFA Immunization and Active EAE Induction

Mice were immunized subcutaneously at two sites of the chest (50 µl/side) with an emulsion containing 100 μg MOG p35-55 (Genemed Synthesis) and CFA containing 200 μg of heat-killed *Mycobacterium tuberculosis* (H37RA, Difco Laboratories) [43]. For active EAE induction, mice were also injected with 100-250 ng (lot-dependent) *Bordetella pertussis* toxin (PTX, #181, List Biologicals) on days 0 and 2 post-immunization. Mice were examined daily for clinical signs of EAE using a five-point scale: (0) no clinical signs; (1) limp tail; (2) hindlimb or foot weakness; (3) complete hindlimb paralysis in one or both hindlimbs; (4) hindlimb paralysis plus some forelimb weakness; (5) moribund or dead. Mice that died from EAE were assigned a score of five for the remainder of the experiment. Cumulative score was calculated for each mouse by adding up the daily scores over the period of observation [43]. Mice were checked for the hindlimb clasping reflex after tail suspension and it was noted whether the hindlimbs curled inwards (clasping) or remained splayed [30].

### 4.8. Measuring MOG p35-55 Reactive T Cell Recall Assays in Spleen and Draining Lymph Nodes

Mice were immunized with MOG p35-55/CFA, and at day 9 post-immunization, the recall proliferative and cytokine response of the spleen and draining lymph nodes to MOG p35-55 were examined as described previously [43]. In brief, mononuclear cells were isolated from the spleen and draining lymph nodes and cultured in 96-well flat-bottomed plates (0.5 x 10^6^ cells/well) with 0, 2, 5, and 10 μg/ml MOG p35-55 in Complete RPMI Medium (CM): RPMI 1640, 2 mM L-glutamine, 1 mM sodium pyruvate, 0.1 mM nonessential amino acids, 100 U/mL of penicillin, 0.1 mg/ml of streptomycin (all from Life Technologies), 50 µM 2-mercaptoethanol (BioShop), and 10% FBS. Cytokines were measured in culture supernatants at the time of peak production (IFN-γ at 48 h, IL-17A at 72 h) using Ready-Set-Go ELISA kits (ThermoFisher) or the LegendPlex Th cytokine panel (Biolegend). In addition, the number of MOG p35-55 CD4^+^ T cells that had expanded after immunization was estimated in the spleen by examining the number of CD4^+^ T cells that divided as assessed by CFSE dilution assay (Molecular Probes) after re-stimulation *ex vivo* with MOG p35-55. For this assay, 10 million spleen cells were labelled with 0.5 µM CFSE according to the kit instructions. Cells were then plated in duplicate in 24-well plates (2.5 x 10^6^ cells/well) with MOG p35-55 (5 µg/mL) for 72 h in a 37°C incubator with 5% CO_2_. Cells were then harvested, washed with FACS buffer, and stained with LIVE/DEAD™ Fixable Aqua Dead Cell Stain (1:1000) and the Ab cocktail detailed in Suppl. Table 2. The number of MOG p35-55 reactive Th cells was estimated as the frequency of CD44^hi^CD4^+^ T cells that specifically divided in response to MOG p35-55, multiplied by the total amount of live splenocytes counted at the hemacytometer.

### 4.9. Passive transfer EAE

Passive transfer studies were carried out similar to described previously [89]. Eight-week-old female C57BL6/J mice or CD45.1 Pepboy mice were immunized with MOG p35-55/CFA, and mononuclear cells were isolated from spleen and draining lymph nodes at day 9 post-immunization. Cells were plated at 8 x 10^6^/ml in 75 cm^2^ vented tissue culture flasks with 10 µg/ml of MOG p35-55 and 10 ng/mL IL-23 (R&D Systems) for 3 days. Cells were then washed three times in 1x PBS and then 10-20 x 10^6^ cells (experiment-dependent) were injected into recipient AA- or CS-exposed male or female CD45.2 C57BL6/J mice. Exposures to CS and AA were continued throughout the experiment.

### 4.10. Histological Scoring of EAE

Brains and spinal cords from mice with active EAE were isolated, preserved in 10% neutral buffered formalin (Sigma) for 7 days, processed in paraffin, and embedded in a single paraffin block (10–12 spinal cord and 6 brain sections/block) according to our published protocol [88]. Cross-sections (5 μm) were cut and stained with Luxol Fast Blue (LFB) to visualize myelin or CD45 IHC [88]. Percent demyelination and CD45^+^ leukocytes/mm^2^ in white matter were measured using ImageJ [88].

### 4.11. CNS Mononuclear Cell Isolation

CNS mononuclear cells were isolated as described previously using collagenase digestion followed by Percoll gradient[43]. For cytokine measurement, isolated CNS or lung mononuclear cells were incubated with or without 20 µg/ml MOG p35-55 overnight (12 h) in a 37°C incubator with 5% CO_2_. GolgiStop (BD Biosciences) was added in the last 6 h of culture. Cells were washed with FACS buffer and stained for FACS as described above.

### 4.12. Analysis of a human lung tissue microarray dataset

To identify immune response pathways enriched in smoking patients, we leveraged a previously published microarray-derived gene list identifying probe sets significantly altered between smoking and non-smoking patients with lung cancer, obtained from Bossé *et al*. [35]. While raw data is publically available in the Gene Expression Omnibus database (GSE23546), files were not annotated with patient information, and so key parameters required to perform our analyses, such as sex and smoking status, were not available to us, so analysis was performed on the published list of DEGs. All data were analyzed on RStudio (version 4.4.0). GO pathway enrichment analysis was performed using the clusterProfiler R package. Gene symbols were matched to corresponding Entrez Gene identifiers using the org.Hs.eg.db annotation database. Statistical significance was determined using a hypergeometric test with Benjamini–Hochberg correction for multiple hypothesis testing. GO terms were considered enriched at a false discovery rate (FDR) < 0.05. Redundant GO terms were reduced by grouping highly overlapping pathways and selecting representative immune-related biological processes for visualization.

Enrichment of MSigDB Hallmark and Reactome gene sets was assessed using the fgsea algorithm applied to the ranked gene list. We obtained curated pathway collections from the Molecular Signatures Database (MSigDB), using the msigdbr R package, which included Hallmark and Reactome pathway gene sets. Hallmark gene sets represent curated, non-redundant biological processes, while Reactome provides detailed signaling pathway annotations. Using the fgsea R package, genes were ranked according to the differential expression statistic derived from smoking compared to non-smoking. The enrichment score for each gene set was calculated using the running-sum statistic across the ranked gene list. Significance was estimated using the adaptive multilevel splitting Monte-Carlo method implemented in fgseaMultilevel, and FDRs were calculated using the Benjamini–Hochberg method. Gene sets were considered significantly enriched at FDR < 0.05. Leading-edge genes contributing to each enriched pathway were extracted from the fgsea results and used for downstream overlap analyses and visualization. Classical GSEA could not be performed, given our inability to access the full dataset. Results were visualized using dot plots generated with the ggplot2 and dplyr R packages. The heatmap of gene expression Z-scores, across enriched GO immune pathways were generated using the pheatmap R package.

### 4.13. Quantification of Cytokines and Chemokines in Lung Homogenates

Lungs were isolated from AA- or CS-exposed mice 9 days after immunization with MOG p35-55/CFA, snap-frozen in liquid nitrogen and stored at -80°C. Lungs were weighed and homogenized in T-PER buffer according to instructions (Thermo Scientific). Homogenates were incubated on ice for 1 h, centrifuged for 10 min at 10,000 x g at 4°C, aliquoted and frozen at -80°C. Total protein was measured in homogenates using a BCA Assay (Pierce). Cytokine and chemokine levels were measured in supernatants by Eve Technologies using the Mouse Cytokine/Chemokine 32-Plex Discovery Assay Array (MD32) panel, or with the individual ELISA kits: amphiregulin (RayBiotech), BAFF and CXCL13 (R&D Systems), IL-23 and IL-33 (Biolegend), and Verikine-HS Mouse IFN-α all subtype ELISA kit (PBL Biosciences). Levels were expressed per mg of lung protein.

### 4.14. DC: T Cell Co-culture Assays

Mononuclear cells were isolated from pooled lungs of AA- and CS-exposed male C57BL/6J mice (n=5/group) as described above, and DCs were enriched from lung mononuclear cells using the mouse Pan Dendritic Cell Isolation Kit (Miltenyi). Isolated DCs (2.5 x 10^4^) were cultured in triplicate in round-bottom plates with 0 or 6 ng/mL LPS and 0-20 µg/mL MOG p35-55 for 6 h at 37°C and 5% CO_2_ (pre-activation phase). Splenic CD4^+^ T cells were enriched from 2D2 male mice using the MojoSort Mouse CD4 T Cell Isolation Kit (Biolegend). A portion of these cells were stained with Cell Proliferation Dye eFluor450 according to the product directions (eBioscience). 2D2 CD4^+^ T cells (5.0 x 10^4^) were placed on top of the cultured DCs. These cells were then cultured for 2-3 days at 37°C and 5% CO_2_, and then washed with FACS buffer. Cell proliferation was assessed by dye dilution in 2D2 CD44^hi^CD4^+^ T cells using flow cytometry. For cytokine assessment, cells were re-stimulated with 50 ng/mL PMA and 500 ng/mL ionomycin for 4 h, with GolgiStop added in the last 2 h of culture, and then stained for FACS as described above.

To assess the role of IL-12p40 signalling in T cell priming by lung DCs, DC:T cell co-cultures were set up similarly, with the addition of 10 µg/mL of either anti-mouse IL-12p40 neutralizing Ab (clone C17.8, BioLegend) or isotype control Ab (clone RTK2758, BioLegend) during the pre-activation phase when DCs were incubated with LPS and MOG p35-55. After the co-culture of DCs and 2D2 CD4^+^ T cells for 2-3 days at 37°C and 5% CO_2_, cells were stained and analyzed as described above.

### 4.15. IL-12p40 Blockade in Lung

We tested the effect of anti-IL-12p40 Ab (clone 2A3; BioXCell) in neutralizing IL-12p40 in the lungs of naive mice, which have detectable IL-12p40 protein at steady-state. This was done by administering 50, 125, or 200 µg/mouse of the InVivoMab anti-mouse IL-12p40 Ab (clone C17.8; BioxCell) to mice (n=2/group) intranasally or intratracheally, and then euthanizing mice after 3 days for measurement of IL-12p40 in lung, spleen, and small intestine homogenates using an IL-12p40 ELISA kit (Biolegend). This study demonstrated that 125 µg of anti-IL-12p40 administered intranasally specifically lessened IL-12p40 levels in the lung, while higher doses, or intratracheal administration, also led to systemic depletion of the cytokine (Supplementary Fig. 8).

EAE was induced in AA or CS-exposed male recipients with pre-primed female MOG p35-55 reactive CD45.1^+^ cells. On days -2 and 1 post-transfer, the recipient C57BL/6J mice were anaesthetized with isoflurane and received 125 µg/mouse of anti-IL-12p40 Ab or InVivoMAb rat IgG2a isotype control (clone 2A3; BioXCell) intranasally. Mice were euthanized on day 4 post-transfer, and lungs were collected for flow cytometry as described above.

### 4.16. Statistical Analysis

Data were analyzed using GraphPad Prism (v10). Data are showed as mean ± SEM. For studies that contained 2 groups, a two-tailed Mann-Whitney U test was used to determine between-group differences. For experiments with a 2 x 2 design (e.g., sex by smoking or smoking by treatment), data were analyzed using a 2-way ANOVA followed by a Bonferroni post-hoc test. P values of p≤0.05 were considered statistically significant.

### 4.16. Data availability

The data that support the findings of this study are available from the corresponding author, upon request.

## 5. Author contributions and Funding Support

N.A-S, A.H., B.C., and S.E.D. conceived and conducted experiments and analyzed data. N.A-S. drafted the manuscript. D.X. conducted gene expression analysis and contributed to the writing of the manuscript. E.P. conducted CS and AA exposures on mice. M.P. and E.P. conducted histological analysis. J.V.R. conducted clinical and histological scoring for some EAE studies. E.L. and W.L.L helped with lung histology studies. T.H. conducted ELISA assays. S.E.D., M.S., and C.R. obtained funding for the project. All authors have reviewed the manuscript.

This work was funded by grants from the National MS society (M.S., C.R., S.E.D.) and MS Canada (S.E.D., C.S., W.L.) and start-up funds from the St. Michael’s Hospital Foundation (S.E.D.).

## 6. Acknowledgements

We thank Dr. Monika Lodyga, Dr. Caterina DiCiano-Oliveira and Dr. Xiaofeng Lu from research facilities at St. Michael’s hospital for help with training personnel in imaging and flow cytometry. We thank staff at the Toronto Center for Phenogenomics for embedding and sectioning of our tissues. We thank the staff of the University Health Network animal facility for their expert care of animals. We thank Dr. Alan Lazarus and Dr. Veronica Miron for helpful discussions and mentorship of trainees.

## 8. Figure Legends

**Supplementary Figure 1. Exposure of mice to CS over eight weeks induces immune cell infiltration in the lung. (**A-D) Sections of lungs of AA- and CS-exposed males were stained for CD45. Arrows show immune cell clusters located close to blood vessels and bronchioles. (B-C) Shown are the number of CD45^+^ cells (B) and the mean area of clusters of CD45^+^ cells in mm^2^ (C). (D-G) Lung mononuclear cells were isolated from AA- and CS-exposed and were analyzed by flow cytometry. (D-E) The total number of live CD45 leukocytes isolated from the lungs. (F) Gating strategy (pre-gated on live singlets) used to identify lung immune cell subsets. (G) Heat map of the numbers of the different immune populations in individual mice (n=5 mice per group). Data in B-E are presented as mean ± SEM. Data are from one representative experiment of at least three that were performed. *: p≤0.05 between groups as determined by two-tailed Mann-Whitney U test (B-E) or 2-way ANOVA followed by Bonferroni post-hoc test (G). In (G), p≤0.05 between male groups is shown with black asterisks, and between female groups with red asterisks.

**Supplementary Figure 2. B and T cells accumulate in perivascular and peri-bronchial clusters in cigarette smoke (CS)-exposed lungs.** Dual CD3 (DAB) and B220 (Vector Blue) staining in lung sections of ambient air (AA) and CS-exposed mice. (A) Presence of scattered T cells in the AA-exposed lung. (B-D) Examples of pathology in the CS-exposed lungs. (B) Shows a larger and smaller lymphocytic infiltrate containing T cells and B cells (arrows). (C) Shows a less organized lymphocytic structure around the airway with T and B cells present (arrows). (D) Shows an example of T cell clusters with no B cells present.

**Supplementary Figure 3. Effect of CS exposure on pMOG-reactive Th cytokine responses in female mice during active EAE.** A-B) Ambient air (AA)- or CS-exposed female C57Bl/6J mice were immunized with MOG_35-55_ to assess the *in vitro* cytokine and proliferative recall responses of cells isolated from spleen and draining lymph nodes (dLNs) 9 days after immunization. Levels of Th1, Th17 and Th2 cytokines in supernatants of MOG_35-55_-stimulated (0-10 µg/mL) cell cultures from spleens (A) and dLNs (B). Data are mean ± SEM of triplicate wells of cultures from pooled mice from one experiment that is representative of two performed. *: p≤0.05 between groups as determined by Mann-Whitney test.

**Supplementary Figure 4. EAE is still attenuated in male mice if CS-exposures are stopped prior to the induction of EAE.** Male C57BL/6J mice were exposed to 8-10 weeks of AA or CS, followed of a 4.5 day washout period before active EAE induction according to the schematic in (A). EAE scores (B) and cumulative EAE scores (C). Data are presented as mean ± SEM of individual mice from one experiment. *: p≤0.05 between groups as determined by two-way ANOVA and Bonferroni post hoc test (B) or by two-tailed Mann-Whitney U test (C).

**Supplementary Figure 5. CS also induces a hyperactivation of female Th cells in passive transfer EAE, but does not activate the host cells.** (A) Passive EAE was induced in male (black) or female (red) C57BL/6J mice that had been exposed to ambient air (AA) and CS, by transferring IL-23-polarized, MOG_35-55_-reactive female T cells from unexposed donor mice. Mononuclear cells were isolated from the lungs and the spleen at a time when EAE was delayed in the CS group. (A) Numbers of MOG_35-55_-reactive T cells producing IL17, co-producing IL17 and IFN-γ, or producing IFN-γ in spleens of male mice during passive transfer EAE as assessed by flow cytometry after stimulation *in vitro* with MOG_35-55_ for 12 hours, with GolgiStop added in the last 6 h. (B-D) Results from an experiment done in females mice. (B) The numberof donor pMOG-reactive T cells detected in the AA- and CS-exposed female lungs in passive EAE. (C) The expressions CD25 and CD69 by median fluorescence intensity (MFI) in donor T cells in the lungs of female recipients. (D) Shows the clinical scores of mice in a companion experiment where females were followed for clinical signs. (E) Shows expressions of CD25 and CD69 in recipient T cells (CD45.1^-^ CD4^+^) in the lungs of an experiment done in males. Data in A and E are mean + SEM of individual mice in one experiment that is representative of 4 independent experiments that were performed. Data points in B-D are individual mice from one experiment of two that were performed. *: p≤0.05 between groups as determined by two-way ANOVA and Bonferroni post hoc test (A) or by two-tailed Mann-Whitney U test (B-D).

**Supplementary Figure 6. Cigarette smoke (CS) enhances cytokine production by 2D2 CD4^+^ T cells in the lungs.** Male 2D2 mice aged 6-7 weeks were exposed to ambient air (AA) or CS. Mice were followed for clinical signs and flow cytometry was performed at endpoint to examine the profile of 2D2 CD4^+^ T cells in the lung and spleen. A) Total number of cells in lungs. B) Total number of myelin-reactive, transgenic TCR-expressing, 2D2 CD4^+^ T cells in the lungs. C) Distribution of naïve (CD44^-^CD62L^+^), central memory (CD44^+^CD62L^+^) and effector memory (CD44^+^CD62L^-^) in myelin-reactive 2D2 CD4^+^ T cells in the lungs in the AA- and CS-exposed mice. D-E) Median fluorescence intensity (MFI) of CD25 (D) and CD69 (E) in memory (CD44^high^) and naïve (CD44^low^) myelin-reactive 2D2 CD4^+^ T cells in the lungs. F) MFI of IFN-γ and IL-17 in lung cytokine-producing, memory 2D2 CD4^+^ T cells cultured *in vitro* with/without MOG_35-55_. G) Representative FACS histograms of IFN-γ and IL-17 production by lung cytokine-producing, memory 2D2 CD4^+^ T cells. G) Total number of cells in spleen. H) Total number of myelin-reactive, transgenic TCR-expressing, 2D2 CD4^+^ T cells in the spleen. I) Distribution of naïve (CD44^-^CD62L^+^), central memory (CD44^+^CD62L^+^) and effector memory (CD44^+^CD62L^-^) phenotypes in myelin-reactive 2D2 CD4^+^ T cells in the spleen. J) Median fluorescence intensity (MFI) of CD69 in memory (CD44^high^) myelin-reactive 2D2 CD4^+^ T cells in the spleen. K-L) MFI of IFN-γ and IL-17 in splenic cytokine-producing, memory 2D2 CD4^+^ T cells cultured *in vitro* with/without MOG_35-55_. Data are presented as mean ± SEM of individual mice (n=8-9/group) from one experiment. *: p≤0.05 between groups as determined by two-tailed Mann-Whitney U test (A-B, C-D, E, G-L), χ2 test (C, I), and two-way ANOVA and Bonferroni post hoc test (F).

**Supplementary Figure 7. Cigarette smoke (CS) increases the activation of antigen presenting cells (APCs) in the lung.** Female C57BL/6J mice were exposed to ambient air (AA) or CS for 8-10 weeks and then were transferred with MOG p35-55 reactive T cells. Lungs were harvested at a time when AA-exposed mice had developed EAE and CS-exposed mice were still asymptomatic. Representative FACS plots of the expression of activation markers (CD86, MHC-II) in APCs isolated from female C57BL/6J mice exposed to AA or CS.

**Supplementary Figure 8. Blockade of lung IL-12p40 production by intranasal vs intratracheal administration of a blocking anti-IL-12p40 antibody.** (A) Blocking anti-IL12p40 Ab (0-250 µg/animal) was administered either intranasally or intratracheally to male C57BL/6J mice; three days after the administration, IL-12p40 levels were measured in tissue lysates of lungs, small intestine, and spleen using an ELISA kit that used different antibodies from the ones administered. Data are presented as mean ± SEM of individual mice from one experiment *: p≤0.05 between groups as determined by two-way ANOVA and Bonferroni post hoc test; asterisks show statistical significance between 0 and the marked concentration of Ab for intranasal (black asterisks) and intratracheal (grey asterisks) administration.

