## Supplementary Figures for "Cigarette smoke sets up a pro-inflammatory circuit in the lung that induces the hyper-activation of autoreactive T helper cells"

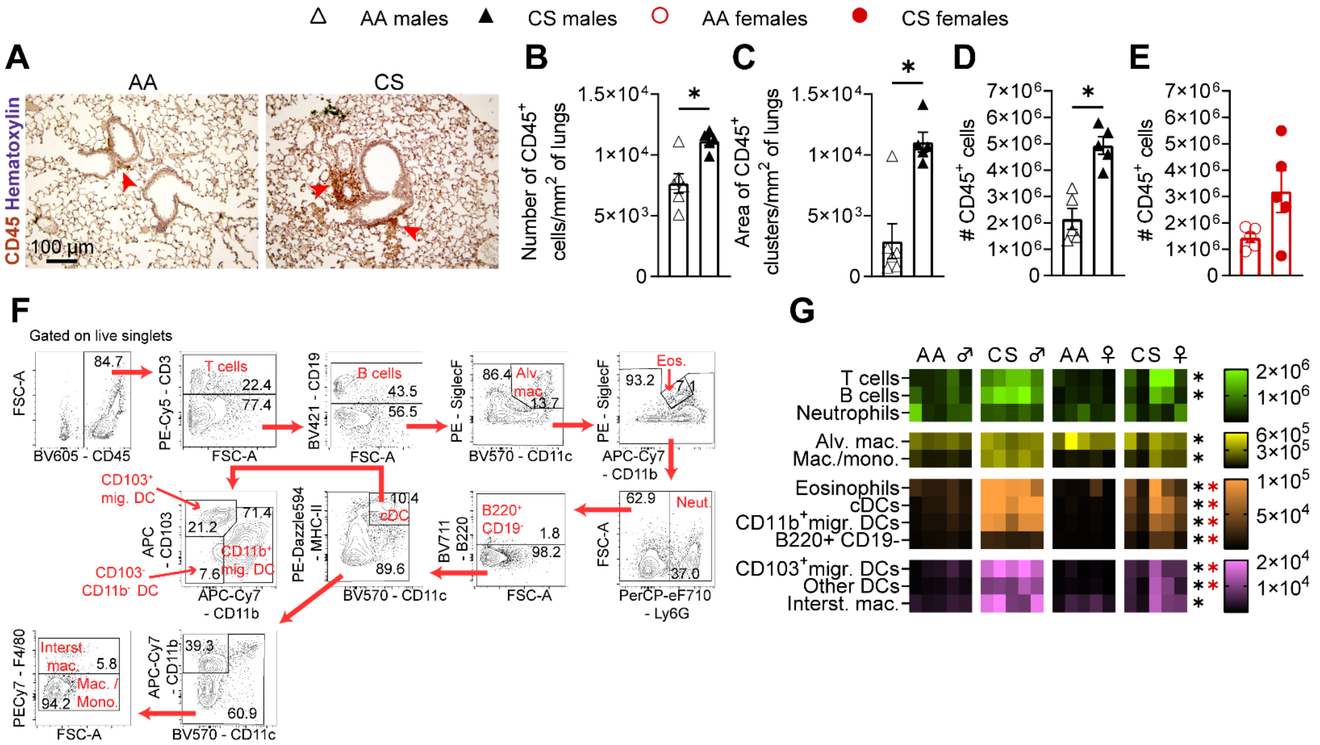


**Supplementary Figure 1. Exposure of mice to CS over eight weeks induces immune cell infiltration in the lung.** (A-D) Sections of lungs of AA- and CS-exposed males were stained for CD45. Arrows show immune cell clusters located close to blood vessels and bronchioles. (B-C) Shown are the number of $CD45^+$ cells (B) and the mean area of clusters of $CD45^+$ cells in $mm^2$ (C). (D-G) Lung mononuclear cells were isolated from AA- and CS-exposed and were analyzed by flow cytometry. (D-E) The total number of live CD45 leukocytes isolated from the lungs. (F) Gating strategy (pre-gated on live singlets) used to identify lung immune cell subsets. (G) Heat map of the numbers of the different immune populations in individual mice (n=5 mice per group). Data in B-E are presented as mean ± SEM. Data are from one representative experiment of at least three that were performed. *: $p \leq 0.05$ between groups as determined by two-tailed Mann-Whitney U test (B-E) or 2-way ANOVA followed by Bonferroni post-hoc test (G). In (G), $p \leq 0.05$ between male groups is shown with black asterisks, and between female groups with red asterisks.

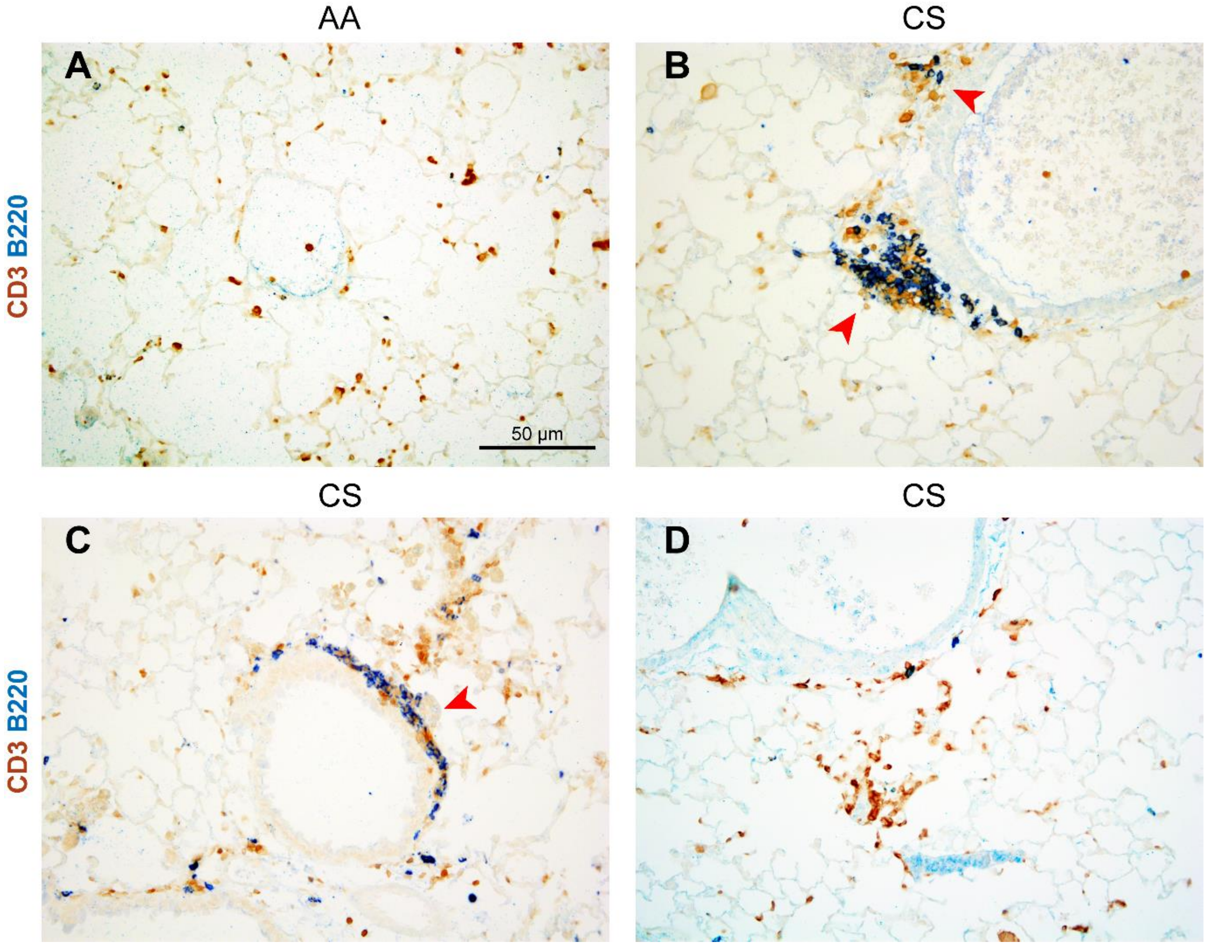


**Supplementary Figure 2. B and T cells accumulate in perivascular and peri-bronchial clusters in cigarette smoke (CS)-exposed lungs.** Dual CD3 (DAB) and B220 (Vector Blue) staining in lung sections of ambient air (AA) and CS-exposed mice. (A) Presence of scattered T cells in the AA-exposed lung. (B-D) Examples of pathology in the CS-exposed lungs. (B) Shows a larger and smaller lymphocytic infiltrate containing T cells and B cells (arrows). (C) Shows a less organized lymphocytic structure around the airway with T and B cells present (arrows). (D) Shows an example of T cell clusters with no B cells present.

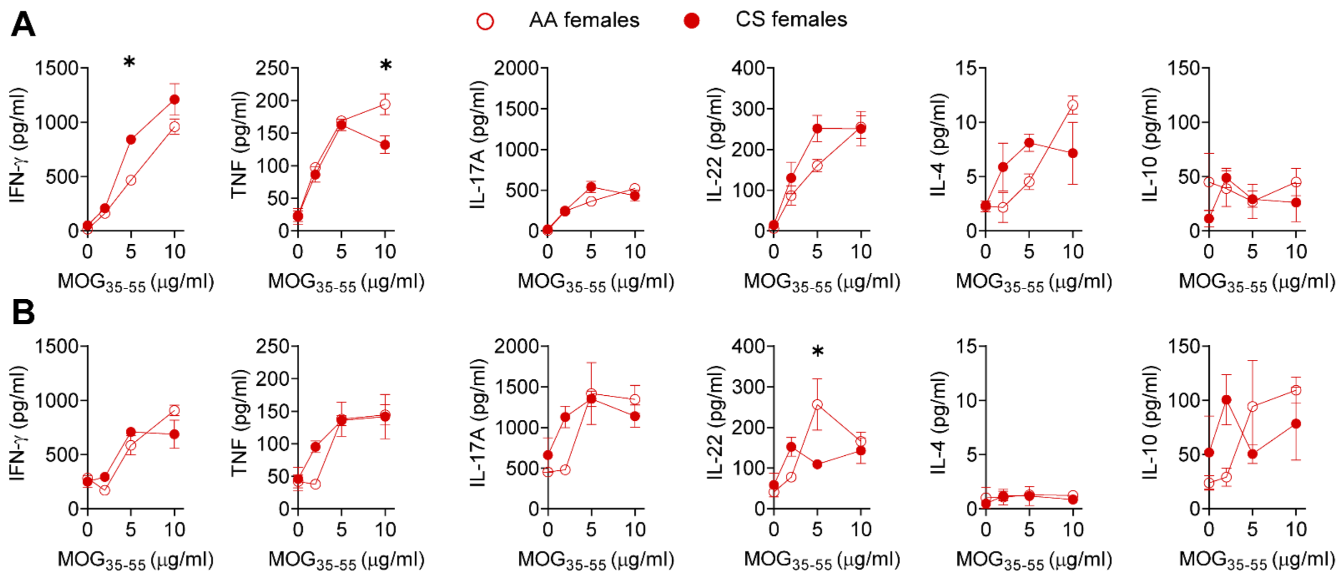


**Supplementary Figure 3. Effect of CS exposure on pMOG-reactive Th cytokine responses in female mice during active EAE.** A-B) Ambient air (AA)- or CS-exposed female C57Bl/6J mice were immunized with $MOG_{35-55}$ to assess the *in vitro* cytokine and proliferative recall responses of cells isolated from spleen and draining lymph nodes (dLNs) 9 days after immunization. Levels of Th1, Th17 and Th2 cytokines in supernatants of $MOG_{35-55}$-stimulated (0-10 μg/mL) cell cultures from spleens (A) and dLNs (B). Data are mean ± SEM of triplicate wells of cultures from pooled mice from one experiment that is representative of two performed. *: p≤0.05 between groups as determined by Mann-Whitney test.

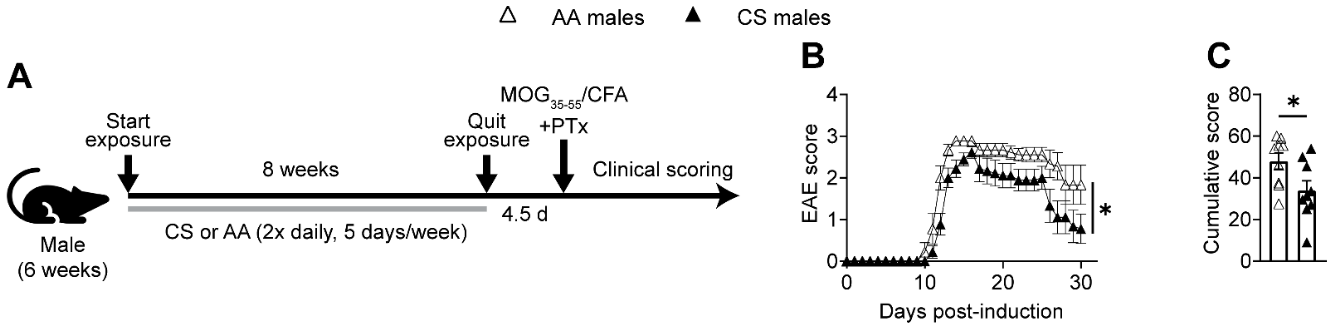


**Supplementary Figure 4. EAE is still attenuated in male mice if CS-exposures are stopped prior to the induction of EAE.** Male C57BL/6J mice were exposed to 8-10 weeks of AA or CS, followed of a 4.5 day washout period before active EAE induction according to the schematic in (A). EAE scores (B) and cumulative EAE scores (C). Data are presented as mean ± SEM of individual mice from one experiment. *: $p \leq 0.05$ between groups as determined by two-way ANOVA and Bonferroni post hoc test (B) or by two-tailed Mann-Whitney U test (C).

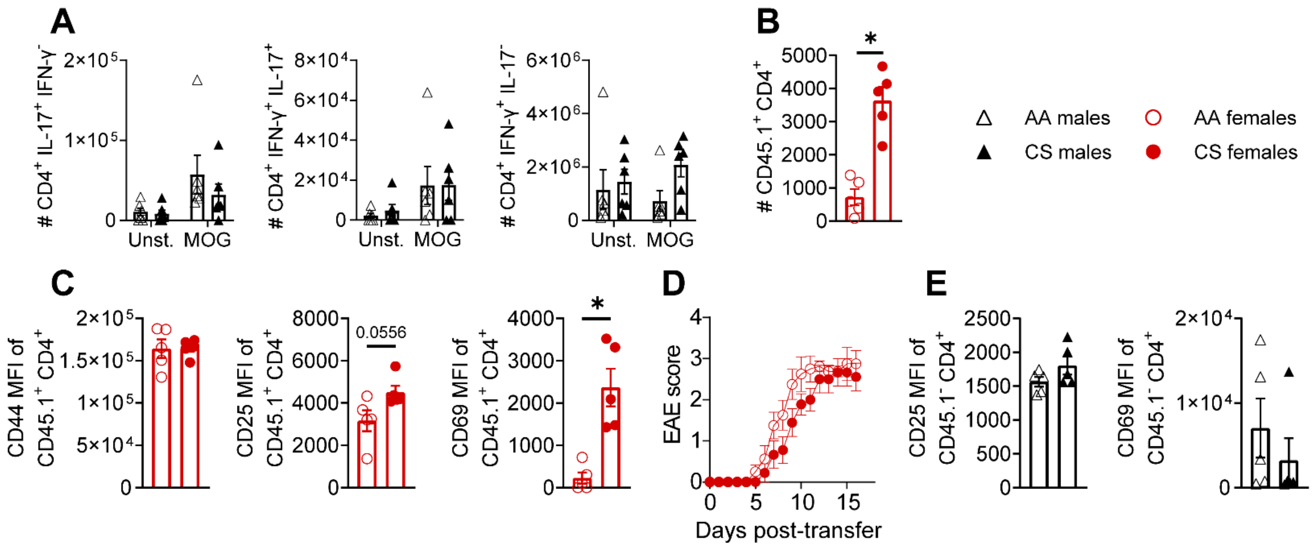


**Supplementary Figure 5. CS also induces a hyperactivation of female Th cells in passive transfer EAE, but does not activate the host cells.** (A) Passive EAE was induced in male (black) or female (red) C57BL/6J mice that had been exposed to ambient air (AA) and CS, by transferring IL-23-polarized, $MOG_{35-55}$-reactive female T cells from unexposed donor mice. Mononuclear cells were isolated from the lungs and the spleen at a time when EAE was delayed in the CS group. (A) Numbers of $MOG_{35-55}$-reactive T cells producing IL17, co-producing IL17 and IFN-γ, or producing IFN-γ in spleens of male mice during passive transfer EAE as assessed by flow cytometry after stimulation *in vitro* with $MOG_{35-55}$ for 12 hours, with GolgiStop added in the last 6 h. (B-D) Results from an experiment done in females mice. (B) The numberof donor pMOG-reactive T cells detected in the AA- and CS-exposed female lungs in passive EAE. (C) The expressions CD25 and CD69 by median fluorescence intensity (MFI) in donor T cells in the lungs of female recipients. (D) Shows the clinical scores of mice in a companion experiment where females were followed for clinical signs. (E) Shows expressions of CD25 and CD69 in recipient T cells ($CD45.1^{-}$ $CD4^{+}$) in the lungs of an experiment done in males. Data in A and E are mean + SEM of individual mice in one experiment that is representative of 4 independent experiments that were performed. Data points in B-D are individual mice from one experiment of two that were performed. *:

$p \leq 0.05$ between groups as determined by two-way ANOVA and Bonferroni post hoc test (A) or by two-tailed Mann-Whitney U test (B-D).

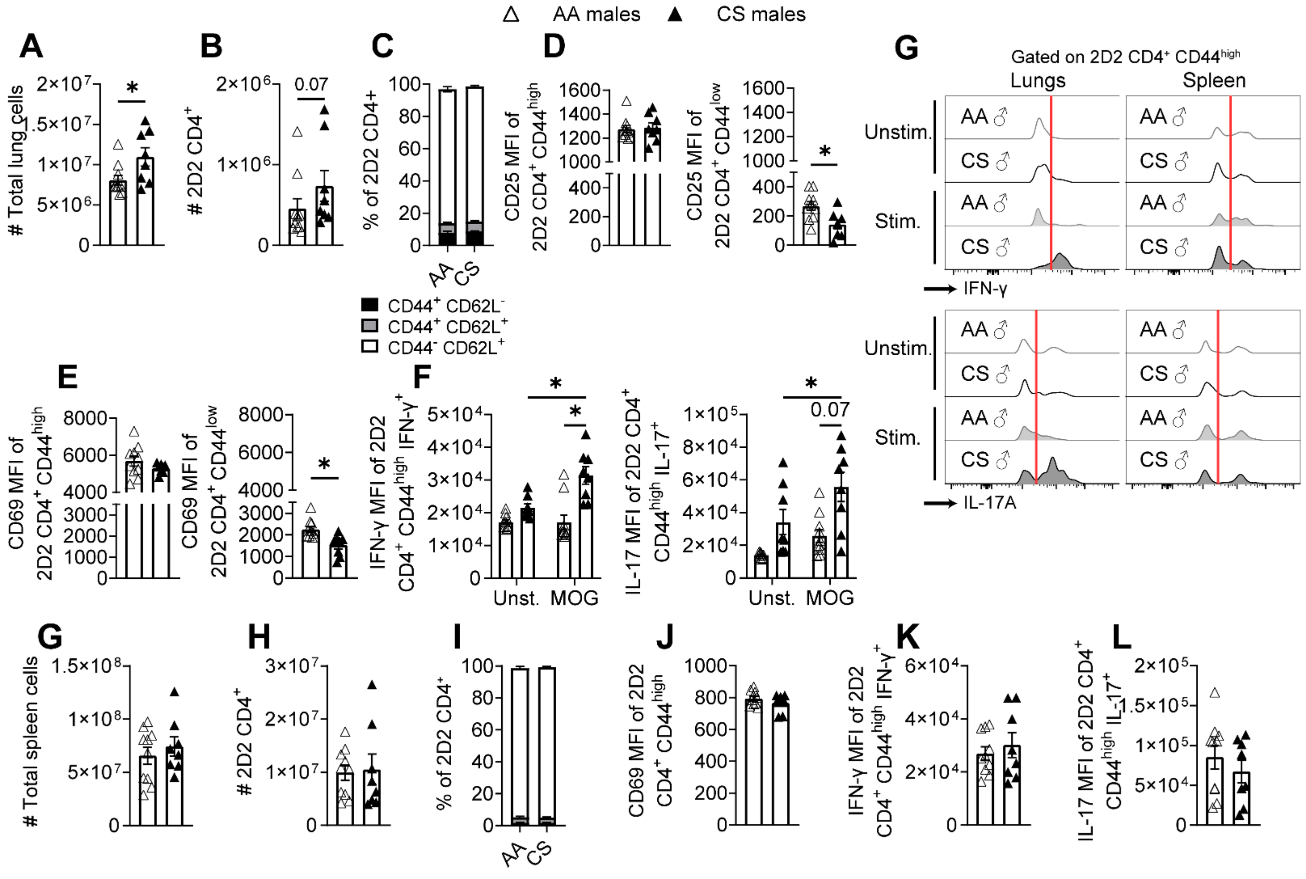


**Supplementary Figure 6. Cigarette smoke (CS) enhances cytokine production by 2D2 CD4⁺ T cells in the lungs.** Male 2D2 mice aged 6-7 weeks were exposed to ambient air (AA) or CS. Mice were followed for clinical signs and flow cytometry was performed at endpoint to examine the profile of 2D2 CD4⁺ T cells in the lung and spleen. A) Total number of cells in lungs. B) Total number of myelin-reactive, transgenic TCR-expressing, 2D2 CD4⁺ T cells in the lungs. C) Distribution of naïve (CD44⁻ CD62L⁺), central memory (CD44⁺CD62L⁺) and effector memory (CD44⁺CD62L⁻) in myelin-reactive 2D2 CD4⁺ T cells in the lungs in the AA- and CS-exposed mice. D-E) Median fluorescence intensity (MFI) of CD25 (D) and CD69 (E) in memory ($CD44^{high}$) and naïve ($CD44^{low}$) myelin-reactive 2D2 CD4⁺ T cells in the lungs. F) MFI of IFN-γ and IL-17 in lung cytokine-producing, memory 2D2 CD4⁺ T cells cultured *in vitro* with/without $MOG_{35-55}$. G) Representative FACS histograms of IFN-γ and IL-17 production by lung cytokine-producing, memory 2D2 CD4⁺ T cells. G) Total number of cells in spleen.

H) Total number of myelin-reactive, transgenic TCR-expressing, 2D2 $CD4^+$ T cells in the spleen. I) Distribution of naïve ($CD44^-CD62L^+$), central memory ($CD44^+CD62L^+$) and effector memory ($CD44^+CD62L^-$) phenotypes in myelin-reactive 2D2 $CD4^+$ T cells in the spleen. J) Median fluorescence intensity (MFI) of CD69 in memory ($CD44^{high}$) myelin-reactive 2D2 $CD4^+$ T cells in the spleen. K-L) MFI of IFN-γ and IL-17 in splenic cytokine-producing, memory 2D2 $CD4^+$ T cells cultured *in vitro* with/without $MOG_{35-55}$. Data are presented as mean ± SEM of individual mice (n=8-9/group) from one experiment. *: $p \leq 0.05$ between groups as determined by two-tailed Mann-Whitney U test (A-B, C-D, E, G-L), $\chi 2$ test (C, I), and two-way ANOVA and Bonferroni post hoc test (F).

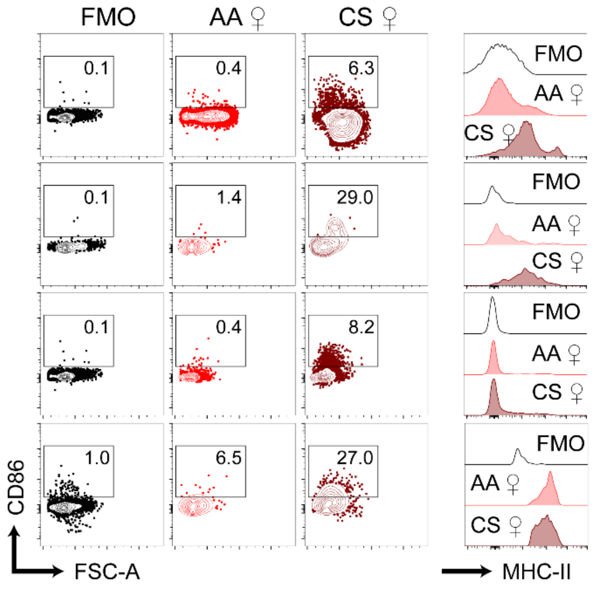


**Supplementary Figure 7. Cigarette smoke (CS) increases the activation of antigen presenting cells (APCs) in the lung.** Female C57BL/6J mice were exposed to ambient air (AA) or CS for 8-10 weeks and then were transferred with MOG p35-55 reactive T cells. Lungs were harvested at a time when AA-exposed mice had developed EAE and CS-exposed mice were still asymptomatic. Representative FACS plots of the expression of activation markers (CD86, MHC-II) in APCs isolated from female C57BL/6J mice exposed to AA or CS.

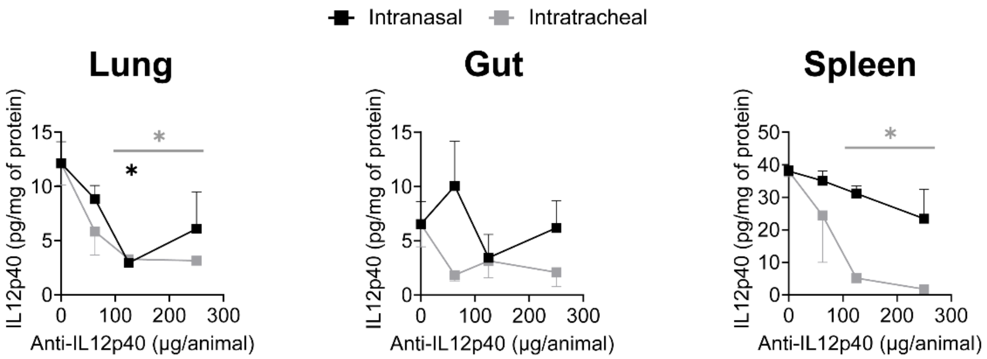


**Supplementary Figure 8. Blockade of lung IL-12p40 production by intranasal vs intratracheal administration of a blocking anti-IL-12p40 antibody.** (A) Blocking anti-IL12p40 Ab (0-250 μg/animal) was administered either intranasally or intratracheally to male C57BL/6J mice; three days after the administration, IL-12p40 levels were measured in tissue lysates of lungs, small intestine, and spleen using an ELISA kit that used different antibodies from the ones administered. Data are presented as mean ± SEM of individual mice from one experiment *: $p \leq 0.05$ between groups as determined by two-way ANOVA and Bonferroni post hoc test; asterisks show statistical significance between 0 and the marked concentration of Ab for intranasal (black asterisks) and intratracheal (grey asterisks) administration.
