## Supplementary Table 2 for "Cigarette smoke sets up a pro-inflammatory circuit in the lung that induces the hyper-activation of autoreactive T helper cells"

**Supplementary Table 2.** Antibodies used for FACS, organized by panel.

| Panel | Specificity | Fluorochrome | Clone | Vendor | Extracellular /Intracellular |
|---|---|---|---|---|---|
| **T Cell Proliferation (Figure 1C-E)** | CD4 | PE-Cy7 | RM4-5 | Invitrogen | Extracellular |
| | CD44 | PE-Cy5 | IM7 | BioLegend | |
| **Cytokine Production in CD45.1 AT-EAE (Figure 3B-F)** | CD45.1 | PE | A20 | BD Biosciences | Extracellular |
| | CD4 | BV785 | RM4-5 | BioLegend | |
| | CD44 | PE-Cy5 | IM7 | BioLegend | |
| | IFN-γ | BV605 | XMG1.2 | BioLegend | Intracellular |
| | IL-17A | APC | TC11-18H10.1 | BioLegend | |
| | GM-CSF | FITC | MP1-22E9 | Invitrogen | |
| **Cytokine Production in AT-EAE, Panel 1 (Figure 3G-I; Suppl. Figure 5A)** | CD45 | PE-Cy7 | 30-F11 | BioLegend | Extracellular |
| | CD4 | FITC | RM4-4 | BD Biosciences | |
| | IFN-γ | PE | XMG1.2 | Invitrogen | Intracellular |
| | IL-17A | APC | TC11-18H10.1 | BioLegend | |
| **Cytokine Production in AT-EAE, Panel 2 (Figure 3I)** | CD45 | PE-Cy7 | 30-F11 | BioLegend | Extracellular |
| | CD4 | APC-Cy7 | GK1.5 | BioLegend | |
| | IFN-γ | PE | XMG1.2 | Invitrogen | Intracellular |
| | IL-17A | APC | TC11-18H10.1 | BioLegend | |
| | GM-CSF | FITC | MP1-22E9 | Invitrogen | |
| **Activation of T cells (Fig. 3 J; Suppl. Figure 5B-D)** | CD45.1 | FITC | A20 | eBioscience | Extracellular |
| | CD3 | PerCP-Cy5.5 | 17A2 | BioLegend | |
| | CD4 | APC-Cy7 | GK1.5 | BioLegend | |
| | CD8α | AF700 | 53-6.7 | BioLegend | |
| | CD44 | PE-Cy7 | IM7 | BioLegend | |
| | CD69 | PE | H1.2F3 | BD Biosciences | |
| | CD25 | eFluor450 | PC61.5 | Invitrogen | |
| | PD-1 | BV605 | 29F.1A12 | BioLegend | |
| **Activation of APCs (Figure 5A, Suppl. Figure 7)** | CD45 | BV605 | 30-F11 | BioLegend | Extracellular |
| | CD3 | PE-Cy5 | 145-2C11 | BioLegend | |
| | CD11c | BV570 | N418 | BioLegend | |
| | SiglecF | PE | E50-2440 | BD Biosciences | |

| | | | | | |
|---|---|---|---|---|---|
| | CD11b | APC-Cy7 | M1/70 | BioLegend | |
| | Ly6G | PerCP-eFluor710 | 1A8-Ly6g | Invitrogen | |
| | B220 | BV711 | RA3-6B2 | BioLegend | |
| | MHC-II | PE-Dazzle594 | M5/114.15.2 | BioLegend | |
| | CD103 | APC | 2E7 | eBioscience | |
| | F4/80 | PE-Cy7 | BMB | Invitrogen | |
| | CD86 | FITC | GL-1 | BioLegend | |
| **DC:T cell Co-culture Panel 1 (Figure 5B-D)** | CD3 | PerCP-Cy5.5 | 17A2 | BioLegend | Extracellular |
| | CD4 | APC-Cy7 | GK1.5 | BioLegend | |
| | CD44 | PE-Cy7 | IM7 | BioLegend | |
| | CD69 | FITC | H1.2F3 | eBioscience | |
| | CD25 | PE | PC61.5 | Invitrogen | |
| **DC:T cell Co-culture Panel 2 (Figure 5E-F)** | CD4 | BV650 | RM4-5 | BioLegend | Extracellular |
| | CD44 | PE-Cy7 | IM7 | BioLegend | |
| | IFN-γ | FITC | XMG1.2 | BioLegend | Intracellular |
| | IL-17 | PE | TC11-18H10 | BD Biosciences | |
| ***In Vivo* IL-12p40 Blockade (Figure 5G-N)** | CD45.1 | PE | A20 | BD Biosciences | Extracellular |
| | CD4 | BV785 | RM4-5 | BioLegend | |
| | CD44 | PE-Cy7 | IM7 | BioLegend | |
| | IFN-γ | BV605 | XMG1.2 | BioLegend | Intracellular |
| | IL-17A | APC | TC11-18H10.1 | BioLegend | |
| | GM-CSF | FITC | MP1-22E9 | Invitrogen | |
| **Lung Immune Cells (Suppl. Figure 1D-G)** | CD45 | BV605 | 30-F11 | BioLegend | Extracellular |
| | CD3 | PE-Cy5 | 145-2C11 | BioLegend | |
| | CD19 | BV421 | 6D5 | BioLegend | |
| | CD11c | BV570 | N418 | BioLegend | |
| | SiglecF | PE | E50-2440 | BD Biosciences | |
| | CD11b | APC-Cy7 | M1/70 | BioLegend | |
| | Ly6G | PerCP-eFluor710 | 1A8-Ly6g | Invitrogen | |
| | B220 | BV711 | RA3-6B2 | BioLegend | |

| | | | | | |
|---|---|---|---|---|---|
| | MHC-II | PE-Dazzle594 | M5/114.15.2 | BioLegend | |
| | CD103 | APC | 2E7 | eBioscience | |
| | F4/80 | PE-Cy7 | BMB | Invitrogen | |
| **Activation of 2D2 T Cells (Suppl. Figure 6)** | CD3 | PerCP-Cy5.5 | 17A2 | BioLegend | Extracellular |
| | TCRVβ11 | PE | RR3-15 | BioLegend | |
| | CD4 | APC-Cy7 | GK1.5 | BioLegend | |
| | CD8α | AF700 | 53-6.7 | BioLegend | |
| | CD44 | PE-Cy7 | IM7 | BioLegend | |
| | CD62L | APC | MEL-14 | BioLegend | |
| | CD69 | FITC | H1.2F3 | eBioscience | |
| | CD25 | eFluor450 | PC61.5 | Invitrogen | |
| | PD-1 | BV605 | 29F.1A12 | BioLegend | |
| **Activation of APCs (Figure 5A, Suppl. Figure 7)** | CD45 | BV605 | 30-F11 | BioLegend | Extracellular |
| | CD3 | PE-Cy5 | 145-2C11 | BioLegend | |
| | CD11c | BV570 | N418 | BioLegend | |
| | SiglecF | PE | E50-2440 | BD Biosciences | |
| | CD11b | APC-Cy7 | M1/70 | BioLegend | |
| | Ly6G | PerCP-eFluor710 | 1A8-Ly6g | Invitrogen | |
| | B220 | BV711 | RA3-6B2 | BioLegend | |
| | MHC-II | PE-Dazzle594 | M5/114.15.2 | BioLegend | |
| | CD103 | APC | 2E7 | eBioscience | |
| | F4/80 | PE-Cy7 | BMB | Invitrogen | |
| | CD86 | FITC | GL-1 | BioLegend | |
